# Sparse and distributed cortical populations mediate sensorimotor integration

**DOI:** 10.1101/2023.09.21.558857

**Authors:** Ravi Pancholi, Andrew Sun-Yan, Maya Laughton, Simon Peron

## Abstract

Touch information is central to sensorimotor integration, yet little is known about how cortical touch and movement representations interact. Touch- and movement-related activity is present in both somatosensory and motor cortices, making both candidate sites for touch-motor interactions. We studied touch-motor interactions in layer 2/3 of the primary vibrissal somatosensory and motor cortices of behaving mice. Volumetric two-photon calcium imaging revealed robust responses to whisker touch, whisking, and licking in both areas. Touch activity was dominated by a sparse population of broadly tuned neurons responsive to multiple whiskers that exhibited longitudinal stability and disproportionately influenced interareal communication. Movement representations were similarly dominated by sparse, stable, reciprocally projecting populations. In both areas, many broadly tuned touch cells also produced robust licking or whisking responses. These touch-licking and touch-whisking neurons showed distinct dynamics suggestive of specific roles in shaping movement. Cortical touch-motor interactions are thus mediated by specialized populations of highly responsive, broadly tuned neurons.

## INTRODUCTION

Tactile information plays a central role in the production of movement^1,2^. In mammals, touch-and movement-related activity is present in somatosensory^3^ and motor cortices^4–6^. Motor areas send an efference copy of movement commands to somatosensory regions^7^, while information from somatosensory areas^8–11^ and thalamus^12,13^ impinges on motor areas. The coordination of these information streams is crucial to the effective production of movement^14^, with perturbations of both somatosensory and motor areas degrading movement^15^. How do the neuronal elements of these circuits implement sensorimotor integration?

Neurons performing sensorimotor integration should possess several properties. First, they should represent features of the stimulus that are important for movement generation and modulation. In somatosensory cortices, a subset of superficial L2/3 neurons that exhibit broader tuning^16–18^ can decode more abstract, movement-relevant somatosensory features, making them well-suited for sensorimotor integration. Second, populations involved in sensorimotor integration should exhibit long term stability. Neurons in both sensory^3,19,20^ and motor^21^ cortices exhibit a basal level of turnover, or ‘representational drift’. Drift poses a challenge for reproducible behavior^22^, and sensorimotor interactions implemented would be facilitated by stable sensory and motor responses. Finally, neurons involved in sensorimotor integration are expected to send information to other cortical areas to coordinate the production of movement^23,24^. An ideal population of neurons for performing sensorimotor integration would thus be tuned to abstract tactile features, exhibit long-term stability, and participate in interareal communication.

Given the relative sparsity of tactile and motor populations in somatosensory^23^ and motor^5^ cortices, the likelihood that these populations interact directly is low. But this is arguably the most straightforward way that somatosensory activity could influence movement – by directly driving motor activity^25^. Indeed, cortical motor areas exhibit increasing activity prior to movement initiation that may reflect evidence integration^26–28^. What is the source of this increasing activity? Cortical excitatory circuitry is organized into assemblies^29^ that exhibit high levels of recurrence^30^, with neurons that represent the same sensory input^31–33^ exhibiting elevated connectivity, enabling pattern completion^34–36^ and amplification^37^. Consequently, even small additions of spikes can bias ongoing activity^38^, movement^39^, and perception^40,41^. If the activation of a sensory assembly activates mixed selectivity neurons that also participate in a specific motor assembly, this should increase the likelihood of the encoded movement. Both somatosensory^42,43^ and motor^5,44^ cortices contain neurons with mixed touch-movement selectivity, raising the question of whether these neurons function to bridge somatosensory and motor assembly activity.

In the mouse whisker system, whisker movement (‘whisking’) and whisker contact (‘touch’) information is present in the primary vibrissal somatosensory cortex (vS1)^3,37,45^, while whisking, touch, and licking information are present in the primary vibrissal motor cortex (vM1)^5,6^. Vibrissal S1 and vM1 are densely interconnected^46–48^, with vS1 sending touch signals to vM1^9–11^, and vM1 sending touch, whisking, and licking signals to vS1^7^. Both vM1 and vS1 microstimulation can drive whisker movement^49,50^, and vM1 shows licking-related preparatory activity^51^. Both vM1^5,6^ and vS1^42,43^ contain neurons with mixed touch-motor selectivity. These neurons may therefore mediate interactions between touch and motor representations. If this population also exhibits long term stability and is involved in vM1-vS1 interactions, it would be ideally suited to perform sensorimotor integration. We study sensorimotor integration in L2/3 of both vM1 and vS1 using volumetric calcium imaging to densely sample activity in thousands of neurons per animal^52^. This approach reliably captures activity among mixed selectivity neurons. We first characterize whisker touch, whisking, and licking populations in both vS1 and vM1. Next, we ask what distinguishes the neurons that participate in vS1-vM1 interactions by injecting a fluorophore-expressing retrograde AAV^53^ into one area and imaging the other. We then image activity over several weeks to determine whether these different populations contain neurons that exhibit elevated longitudinal stability. Finally, we ask what characterizes mixed selectivity neurons participating in both touch and motor representations. Our results reveal a specialized population of neurons that bridge tactile and motor representations, exhibit elevated long-term stability, and contribute disproportionately to interareal communication.

## RESULTS

### Vibrissal S1 and vM1 contain a diverse mix of touch, whisking, and lick cells

To assess sensorimotor responses throughout both the primary vibrissal somatosensory cortex (vS1) and primary vibrissal motor cortex (vM1), we imaged separate cohorts of mice (**Table S1**) expressing GCaMP6s in all excitatory neurons (Ai162 X Slc17a7-Cre)^54^. Mice were first implanted with a cranial window over either vS1 or vM1 and, following surgical recovery, trimmed to two whiskers (C2, C3; Methods). Mice were then water restricted and trained on an object detection task in which a pole appeared either near the spared whiskers or out of whisking reach (**Figure 1A**). Following a delay, a sound prompted the animal to respond at one of two lickports, with right licks rewarded for pole in-reach trials and left licks rewarded for out-of-reach trials.

**Figure 1.**
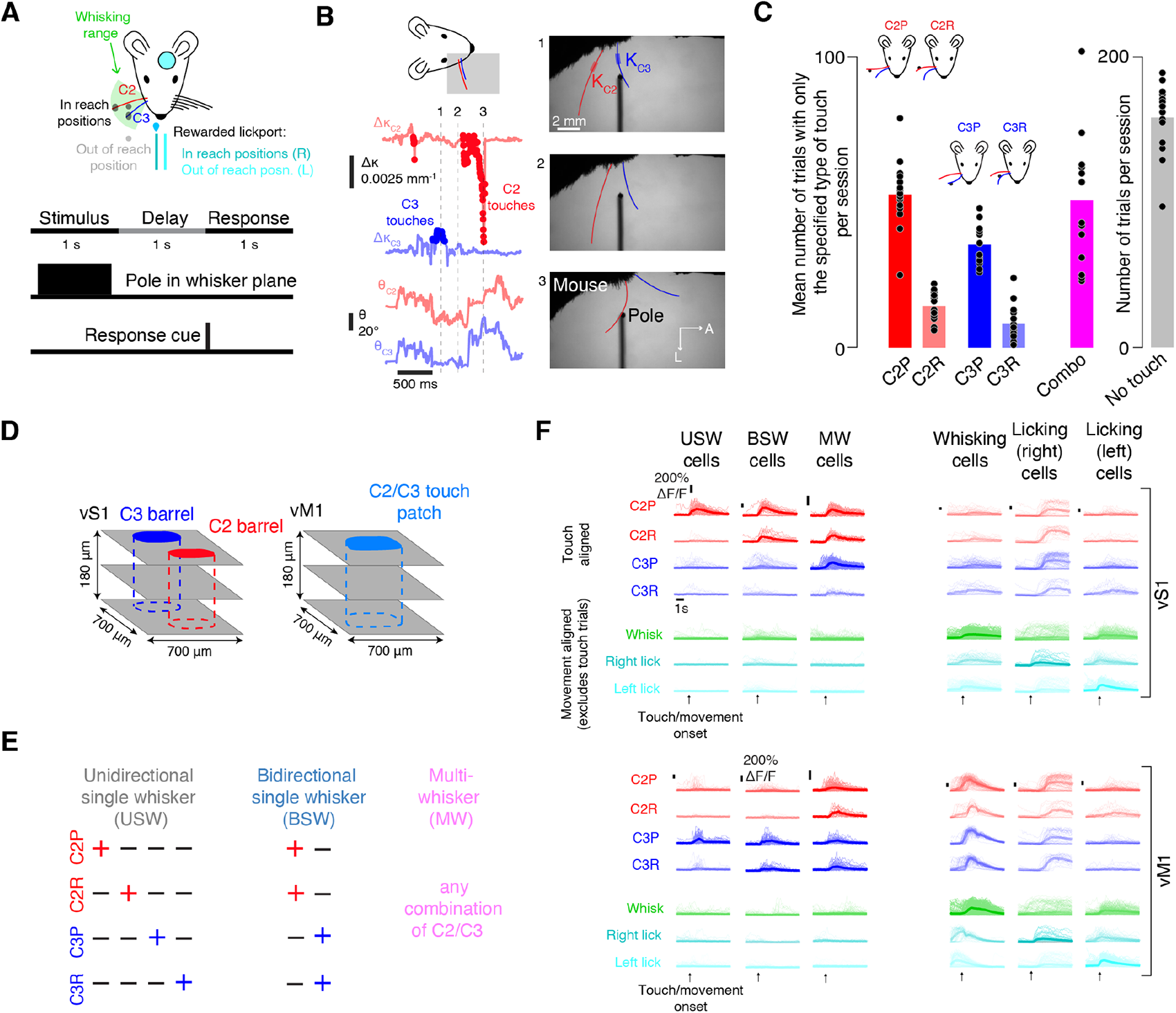
Classification of vS1 and vM1 sensorimotor cell types. (A) Behavioral task. Top, pole appears near the tip of the C2 or C3 whisker or between these two whiskers (black circles) within the whisking range (green), or outside whisking range (grey). The right lickport is rewarded on in reach trials (dark cyan), the left (light cyan) on out of reach trials. Bottom, each trial is composed of a 1 s stimulus epoch where the pole is in the plane of the animal’s whiskers, a 1 s delay epoch, and a 1 s response epoch. (B) Extraction of whisker kinematics. Right, three example frames from a single trial. Thick red lines: whisker segment used for curvature measurements. Left top, curvature change (Δκ) during the example trial; touch times in darker color. Left bottom, angular position of each whisker. (C) Number of trials by touch type (n=16 mice) on first imaging session. C2P, trials with only C2 protractions; C2R, only C2 retractions, etc. ‘Combo’ trials (purple) contain at least two different basic touch types. (D) Imaging strategy. Left, in vS1, imaging was centered on the barrels of the C2 and C3 whiskers. Right, in vM1, imaging was centered on the C2/C3 responsive touch patch. (E) Touch cell classification. Left, unidirectional single whisker cells respond to one of the four basic touch types. Middle, bidirectional single whisker cells respond to both touch directions for one whisker. Right, multiwhisker cells respond to any combination of C2 and C3 touches. (F) Example neurons from a single session. Left, touch responsive neurons. Right, movement-related neurons. For each neuron, response to trials with only one touch type (C2 protractions, C2 retractions, C3 protractions, or C3 retractions) are shown, along with response to movement onset on trials with no touch (whisking onset, right and left licking onset). Top, vS1 example neurons, one per type; bottom, vM1 example neurons. Thick line, mean response; thin lines, individual trial responses. Trial types for which a cell was considered non-responsive are lightened.

Whisker-object touches and whisker movements were tracked using high-speed videography (400 Hz; Methods; **Figure 1B**)^55^. Touches were classified based on the touching whisker’s identity and the direction of touch (**Figure 1C**): C2 protractions (C2P), C2 retractions (C2R), C3 protractions (C3P), and C3 retractions (C3R). Mice produced many trials with isolated touches of only one type on every session (C2P: 52 ± 11 isolated touch trials per session, mean ± SD, n=16 mice; C2R: 14 ± 5; C3P: 35 ± 7; C3R: 8 ± 6), along with trials that involved combinations of these four touch types (51 ± 22 trials per session) and trials with no touch (158 ± 22 trials per session).

Two-photon volumetric calcium imaging^52^ was used to record excitatory neuron activity in each mouse across three 700-by-700 μm planes spaced 60 μm apart in depth (**Figure 1D**), starting at the layer (L) 1-L2 interface and continuing through most of L2/3. For vS1, we imaged the region including the two spared whiskers’ barrels. For vM1, we identified the ‘touch patch’ that contained robust C2 and C3 responses (**Figure S1**). This yielded 2,514 ± 194 vS1 neurons (n=7 mice) or 2,735 ± 681 vM1 neurons (n=9 mice) per animal. Neurons were considered responsive to specific touch types if their mean touch-evoked ΔF/F response exceeded 10% on a given session (Methods). Touch neurons were classified into three categories^16^ (**Figure 1E**): ‘unidirectional single whisker’ (USW) cells responded to only one touch type, ‘bidirectional single whisker’ (BSW) cells responded to both directions for one whisker, and ‘multiwhisker’ (MW) cells responded to at least one touch type for both whiskers. Neurons were classified as ‘whisking’ neurons if their ΔF/F response exceeded 10% following whisking onset in the absence of touch (Methods). Licking neurons were similarly classified and subdivided into left and right lick preferring based on which lick direction they responded to most strongly. We found examples of all three touch cell types, whisking cells, and both lick cell types in vM1 and vS1 (**Figure 1F**).

### Touch responses are disproportionately concentrated among a sparse population of multiwhisker neurons

In primary^16^ and secondary^18^ vibrissal somatosensory cortices, touch activity is disproportionately confined to a minority (1-2%) of multiwhisker cells. We asked whether this pattern was repeated in vM1. We first examined the distribution of touch responses in the imaged population. In superficial vS1 of mice performing the touch detection task, we measured the ΔF/F response to specific single whisker touches (e.g., C2 or C3 protractions; **Figure 2A**). A minority of neurons responded to touch: 16 ± 3% (n=7 mice; **Figure 2B**) of neurons were classified as unidirectional single whisker, 6 ± 1% were bidirectional single whisker, and 8 ± 2% were multiwhisker, consistent with previous reports^16^.

**Figure 2.**
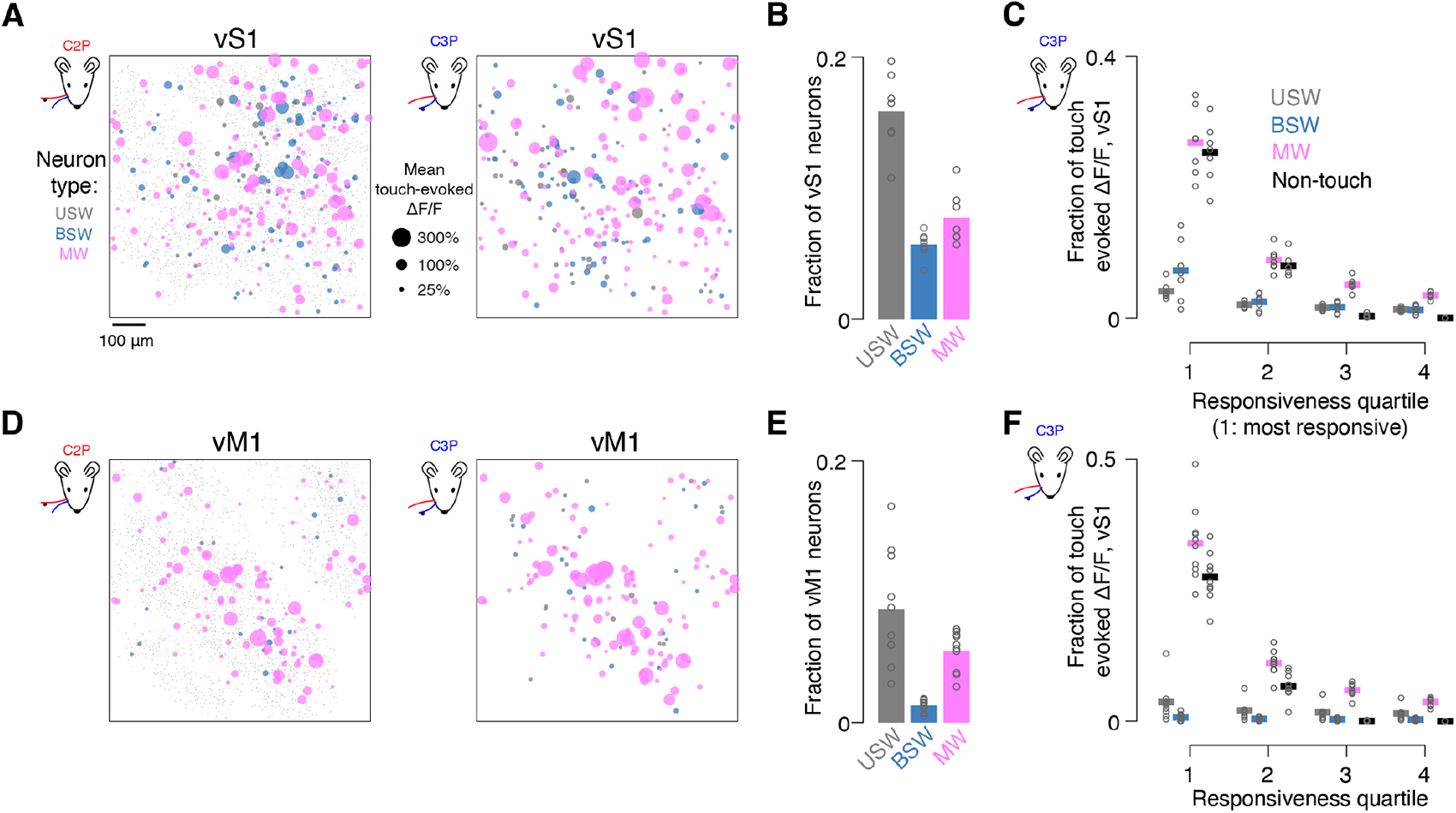
Multiwhisker neurons carry a disproportionate fraction of the vS1 and vM1 touch response. (A) Example vS1 touch responses. Projected view of all three imaged planes in one mouse showing mean touch-evoked ΔF/F across all touch responsive neurons (grey, USW; blue, BSW; magenta, MW) to specific whisker touches (Left, C2P; right, C3P). Nonresponsive neurons are included as small dots in the background of the left panel. (B) Fraction of vS1 neurons belonging to specific whisker representations. Bar, cross-animal mean. Circles, individual animals. (C) Fraction of touch-evoked ΔF/F contributed by each touch neuron type separated into quartiles by responsiveness (Methods). Only neurons that were responsive to C3P touches (Methods) were included. Left, most responsive 25% of neurons of a given neuron type. Thick line, cross-animal mean. Circles, individual animals. For an individual animal, all 16 values shown sum to 1. (D-F) Same as A-C but for vM1.

We next examined touch concentration among multiwhisker neurons. We measured the proportion of the overall ΔF/F response confined to specific touch populations for both C2 and C3 protractions, restricting our analysis to these touch types as they were the most numerous and using only neurons from within a given touch population (USW, BSW, or MW) that were responsive to the touch type being analyzed (C2P, C3P; Methods). Multiwhisker cells produced 44 ± 7% of the protraction touch-evoked ΔF/F response, whereas unidirectional single whisker cells accounted for 9 ± 2% and bidirectional single whisker cells for 13 ± 7% (MW vs. USW, p=0.016, Wilcoxon signed rank test, n=7 mice; MW vs. BSW, p=0.002). Non-touch cells accounted for the remaining 34 ± 6%. Multiwhisker cells thus carried disproportionate fraction of touch-evoked activity in vS1.

Was the touch response further concentrated among a subset of multiwhisker neurons? To address this, we partitioned each touch population into four equal-sized quartiles sorted by touch responsiveness,. We assessed the fraction of the overall ΔF/F response confined to each quartile for both protraction types (i.e., C2 and C3 protractions). We first restricted our analysis to touch neurons that were C3 protraction responsive. We found the most responsive quartile of each population carried substantially more touch activity than the other quartiles (**Figure 2C**). The top quartile of unidirectional single whisker cells produced 4 ± 1% (n=7 mice) of the overall vS1 touch response, the top bidirectional single whisker cells produced 7 ± 5%, and the top quartile of multiwhisker cells produced 27 ± 5%. A minority of multiwhisker cells thus carried a quarter of touch activity. A similar pattern was observed for C2 protractions (**Figure S2**).

In vM1, robust touch activity was confined to a subregion of the imaging field and multiwhisker cells were even more dominant (**Figure 2D**). Multiwhisker cells in vM1 were as numerous as unidirectional single whisker cells, but bidirectional single whisker cells were relatively rare (USW fraction: 9 ± 4%, n=9 mice; BSW: 1 ± 0%; MW: 5 ± 2%). Relative to vS1, both unidirectional single whisker and bidirectional single whisker cells were less frequent in vM1, whereas multiwhisker cells appeared in comparable proportion (**Figure 2E**; vS1 vs. vM1 USW percentage, p=0.003, Wilcoxon rank sum test, n=7 vS1, n=9 vM1; BSW, p<0.001; MW, p=0.055).

Were vM1 touch responses concentrated among the most responsive multiwhisker cells? Mulitwhisker cells were responsible for 55 ± 11% (n=9 mice) of the touch-evoked ΔF/F response in vM1, whereas unidirectional single whisker cells accounted for 9 ± 8% of the response and bidirectional single whisker cells accounted for 2 ± 1%. Thus, in both sensory and motor areas, tactile representations are dominated by multiwhisker neurons. As in vS1, activity was further concentrated among the most responsive quartile of each population (**Figure 2F**; percentage of overall touch response in the most responsive quartile: USW: 4 ± 3%; BSW: 1 ± 1%; MW: 34 ± 7%).

Thus, in vS1 and vM1, touch activity is highly concentrated among a sparse population of multiwhisker neurons.

### Whisking and licking responses are concentrated among a sparse subset of neurons

We next assessed sparsity among whisking and licking representations (**Figure 3A**). Both licking and whisking populations were sparse in vS1, with only a small fraction of neurons responding with each movement: 3 ± 1% (n=7 mice) of neurons responded to whisking onset, 6 ± 2% to right (contralateral) licking onset, and 2 ± 1% to left (ipsilateral) licking onset (**Figure 3B**; restricted to no touch trials, Methods).

**Figure 3.**
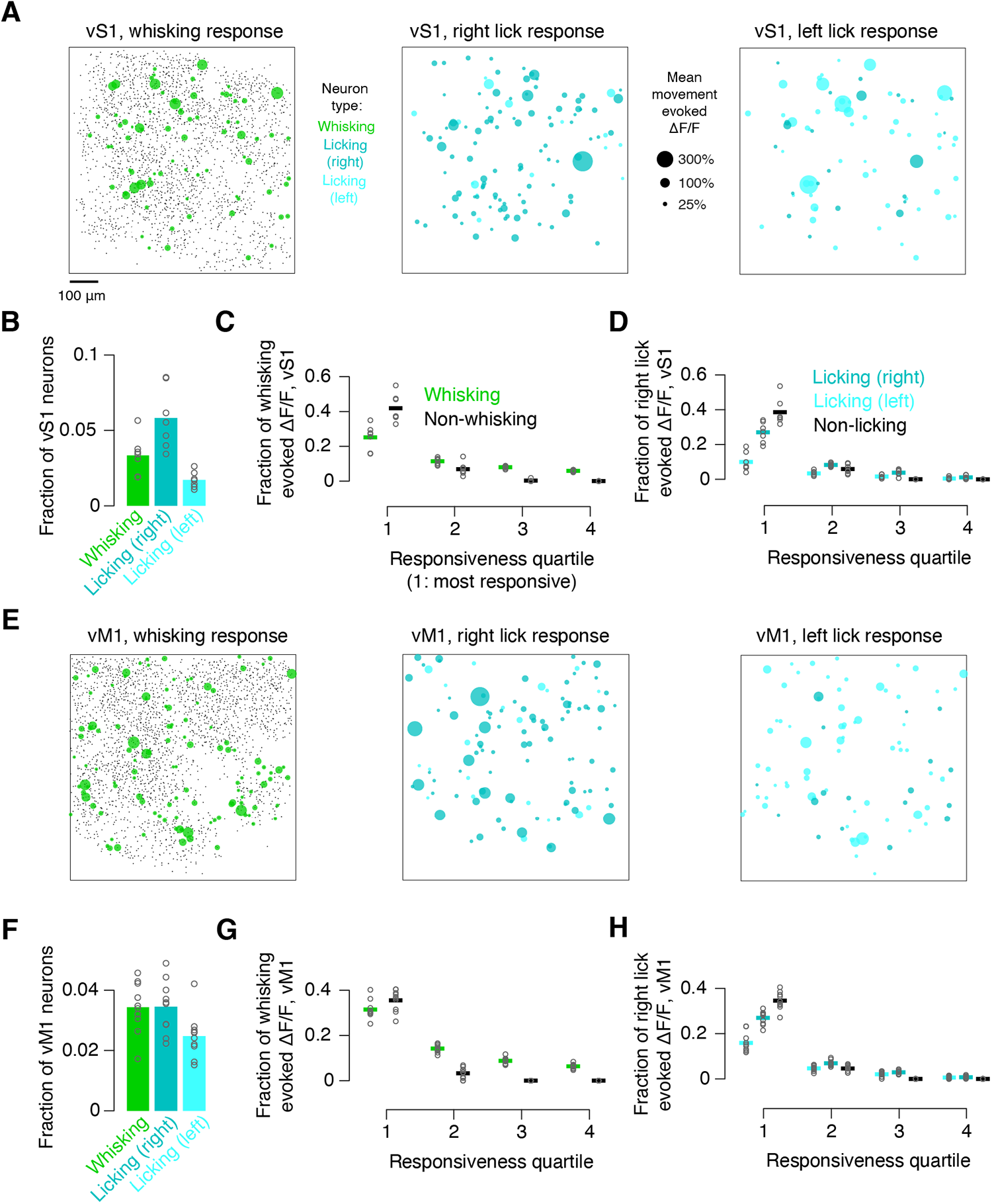
Whisking and licking activity is concentrated in a sparse group of highly responsive neurons. (A) Example vS1 whisking and licking responses. Projected view of all three imaged planes in one mouse showing mean evoked ΔF/F for all responsive neurons Nonresponsive neurons are included as dots in left panel. Left, ΔF/F response following whisking onset. Middle, ΔF/F response following right lick onset (dark cyan, right lick preferring licking cells; light cyan, left lick preferring licking cells). Right, ΔF/F response following left lick onset. (B) Fraction of vS1 neurons belonging to specific movement representations. Bar, cross-animal mean. Circles, individual animals. (C) Fraction of whisking-evoked ΔF/F separated into responsiveness quartiles (Methods). Thick line, cross-animal mean. Circles, individual animals. In all cases, for individual animals, all points sum to 1. (D) As with C, but for fraction of right licking-evoked ΔF/F separated into quartiles by responsiveness (Methods). (E-H) Same as A-D but for vM1.

Whisking cells accounted for 50 ± 10% of vS1 whisking-evoked ΔF/F, with non-whisking cells accounting for 49 ± 10%. For licking, vS1 right licking cells (i.e., contralateral to imaged hemisphere) accounted for 40 ± 7% of the right lick-evoked ΔF/F response, with left licking cells accounting for 15 ± 8% and non-licking cells 44 ± 8%. We next divided neurons of movement representations into quartiles based on their responsiveness and asked what fraction of the overall movement response was confined to the most responsive quartile. Whisking activity was more concentrated among the most responsive whisking neurons (**Figure 3C**), with the top quartile carrying 25 ± 7% of the response. Right-licking neurons in the top responsiveness quartile neurons carried 27 ± 6% of the right lick response (**Figure 3D**). Similar trends were observed for left licks (**Figure S2**). Thus, vS1 whisking and licking responses are highly concentrated.

We next examined movement-related activity in vM1 (**Figure 3E**). Vibrissal M1 licking and whisking populations were sparse (percentage of neurons responding to whisking: 3 ± 1%; right licking: 3 ± 1%; left licking: 2 ± 1%; **Figure 3F**). Overall, vM1 whisking cells accounted for 61 ± 7% of whisking-evoked ΔF/F (non-whisking: 39 ± 7%); for licking, right-lick cells accounted for 38 ± 5% of the right-lick evoked ΔF/F response, and left-lick cells accounted for 23 ± 4% of the right-lick evoked ΔF/F response (non-licking: 39 ± 4%). Whisking responses were further concentrated among the most responsive 25% of a given population (response confined to top quartile: 31 ± 4%; **Figure 3G**). Licking neurons exhibited comparable levels of concentration, with 27 ± 3% of the right lick response confined to the top quartile of right lick neurons (**Figure 3H**; left licks: **Figure S2**).

The fraction of neurons participating in movement-related representations was comparable across vS1 and vM1. Vibrissal M1 had a similar percentage of whisking neurons relative to vS1 (p=0.887, Wilcoxon rank sum test, n=7 vS1, n=9 vM1), a smaller percentage of right lick preferring neurons (p=0.014) and a comparable percentage of left lick preferring neurons (p=0.430). Thus, motor activity is represented in similar fashion across both vS1 and vM1, with a small population of neurons responsible for a large portion of movement-related activity.

### Projections between vS1 and vM1 are dominated by the most responsive neurons

Vibrissal S1 and vM1 are densely interconnected, with neurons encoding touch and whisker movement projecting to vM1^10,11^ and vM1 sending whisking, touch, and licking information to vS1^7^. If the most responsive neurons play a disproportionate role in sensorimotor integration, they should be involved in communication between vS1 and vM1.

We examined the vS1 to vM1 projection by injecting a retrograde AAV^53^ expressing a red fluorophore (rAAV2-retro-FLEX-tdTomato; **Figure S3**) in vM1 and imaging vS1 (**Figure 4A**). We sampled more densely (nine 700-by-700 μm planes spaced 20 μm apart, imaging three planes simultaneously at a time; Methods) to compensate for the rarity of projection neurons. This yielded 5,259 ± 567 vS1 neurons (n=7 mice), of which 296 ± 65 expressed tdTomato (5.6 ± 1.0%, ‘projecting neurons’; Methods). Among touch neurons, multiwhisker neurons projected more frequently than unclassified neurons (i.e., neurons not part of any touch, whisking, or licking population; **Figure 4B**; MW: vs. unclassified: p=0.008, Wilcoxon signed rank test, n=7), whereas unidirectional single whisker neurons projected less frequently than unclassified cells (USW vs. unclassified, p=0.016; BSW vs. unclassified, p=0.219). Both whisking and right-and left-licking neurons projected more than unclassified neurons (whisking vs. unclassified, p=0.031; right licking vs. unclassified, p=0.047; left licking vs. unclassified, p=0.047). Thus, all motor-related populations and multiwhisker touch cells participated disproportionately in the vS1 to vM1 projection.

**Figure 4.**
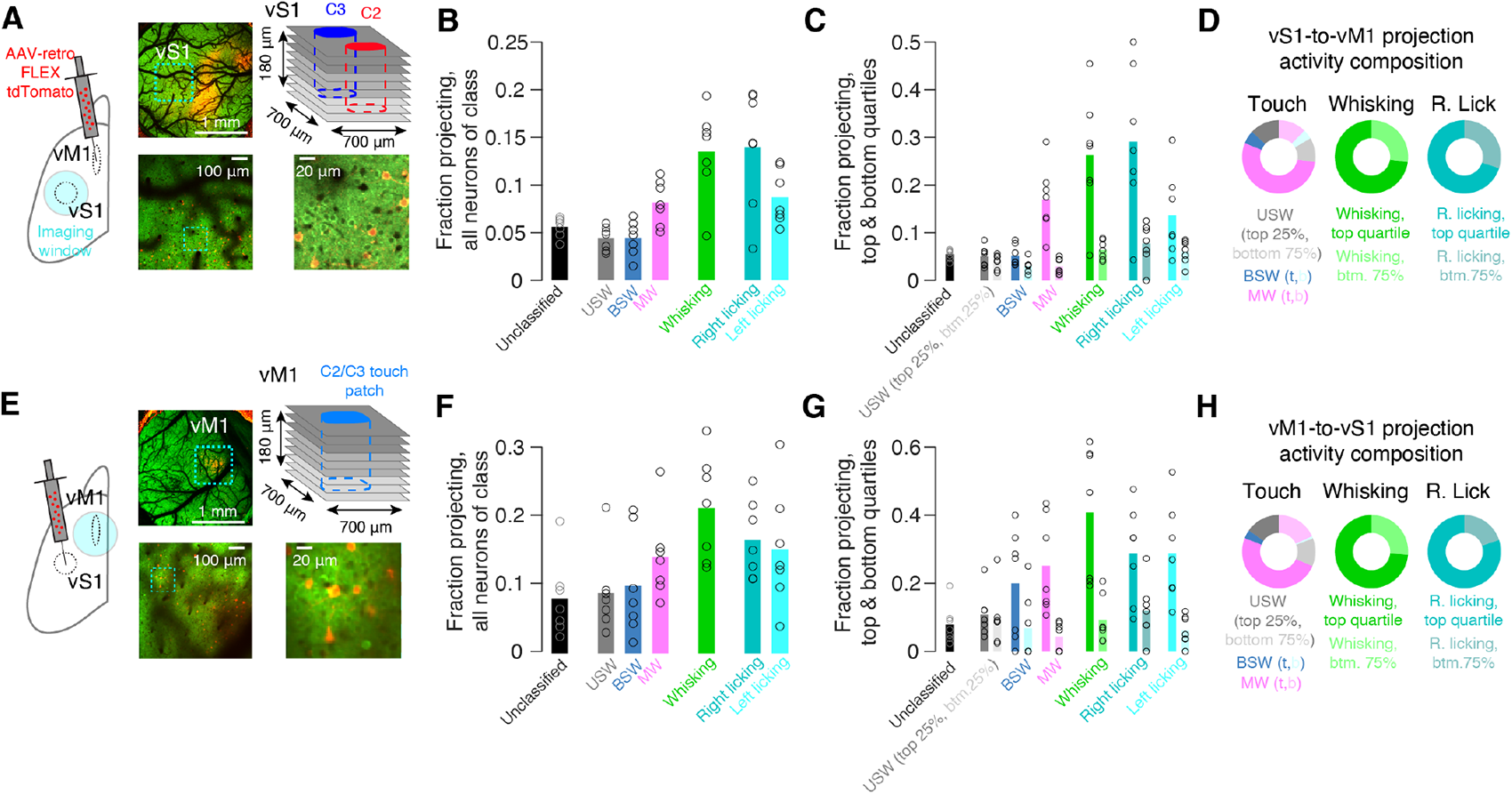
Highly responsive neurons contribute disproportionately to interareal projections. (A) Imaging in vS1 following a retrograde tdTomato virus injection into vM1 (Methods). Left, retrograde injection strategy. Middle top, 4X widefield two-photon image of the cranial window, with inset denoting imaged planes. Middle bottom, example 16X cellular-resolution two-photon imaging plane used to assess individual neuron activity. Right bottom, closer view of a subregion of the imaging plane. Right top, imaging strategy. Planes with same color are imaged simultaneously. (B) Fraction of each type of neuron that expressed red fluorophore. Unclassified neurons did not respond to touch, whisking, or licking. Bars denote mean. Circles, individual mice (n=7). (C) Fraction of each population’s top and bottom responsiveness quartiles that expressed red fluorophore. Darker bars, mean for top quartile; lighter bars, mean for bottom quartile. Circles, individual mice (n=7). (D) Composition of the evoked ΔF/F response to protraction touch, whisking, and right licking carried by specific populations of the projection. Dark color, top quartile. Light color, bottom 75% of responding neurons. (E-H) As in A-D but for imaging of vS1-projecting neurons in vM1.

Did the most responsive neurons project more frequently? Comparing the fraction of neurons from the most responsive quartile of a population that projected to that of the least responsive quartile revealed that the most responsive multiwhisker, whisking, and licking cells were far more likely to project from vS1 to vM1 (**Figure 4C**). Among unidirectional and bidirectional single whisker cells, there was no observed difference (top vs. bottom, USW, p=0.156, Wilcoxon signed rank test; BSW, p=0.078). Among multiwhisker, whisking, and both licking populations, however, the most responsive neurons were far more likely to project than the least responsive neurons (top vs. bottom, MW, p=0.016; whisking, p=0.016; right licking, p=0.031). This same pattern was observed when we analyzed the fraction of total neural activity relayed to the other area (Methods) by the most responsive quartile of each population compared to the least responsive 75%. Among touch neurons, the most responsive quartiles of the three touch types accounted for 53.8 ± 16.8% (MW), 13.1 ± 9.1% (USW), and 5.9 ± 2.9% (BSW) of the projection touch response (**Figure 4D**). Among whisking neurons, the most responsive quartile produced 50.0 ± 13.2% of projection whisking activity. For right licking, the most responsive quartile produced 70.0 ± 15.5% of projection licking activity. The most responsive quartiles of touch, whisking, and licking populations thus contributed disproportionately to vS1-vM1 projection activity.

We next examined the vM1 to vS1 projection (**Figure 4E**). In a separate cohort, we imaged 4,102 ± 775 vM1 neurons (n=7 mice), of which 319 ± 147 (7.7 ± 3.6%) were projecting neurons. Again, multiwhisker, but not unidirectional or bidirectional single whisker touch cells projected with a higher frequency than unclassified neurons (**Figure 4F**; MW vs. unclassified, p=0.016, Wilcoxon signed rank test, n=7; USW vs. unclassified, p=0.219; BSW vs. unclassified, p=0.578). All three movement-related populations projected more frequently than unclassified neurons (whisking vs. unclassified, p=0.016; right licking vs. unclassified, p=0.016; left licking vs. unclassified, p=0.016).

Did the vM1 to vS1 projection disproportionately depend on the most responsive neurons? We found that multiwhisker but not single whisker touch neurons in the top responsiveness quartile were more likely to project than neurons in the bottom quartile (**Figure 4F**; top vs. bottom, Wilcoxon signed rank test, MW, p=0.016; BSW, p=0.156; USW, p=0.812). The most responsive whisking (p=0.016), right-licking (p=0.047), and left-licking (p=0.016) populations also projected more frequently. Projection activity was also concentrated among the most responsive touch cells: the most responsive quartiles of the three touch types accounted for 49.5 ± 12.6% (MW), 15.3 ± 9.0% (USW), and 3.8 ± 3.4% (BSW) of the projection touch response (**Figure 4H**). This was also true for whisking and licking cells, with the most responsive quartile accounting for 48.7 ± 11.4% of projection whisking activity, and the most responsive quartile of right licking neurons accounting for 80.2 ± 15.3% of projection right licking activity.

Thus, between vS1 and vM1, touch activity is primarily relayed by broadly tuned multiwhisker cells and the most responsive multiwhisker touch, whisking, and licking neurons disproportionately contribute to interareal communication.

### The most responsive neurons exhibit elevated longitudinal stability

Sensory^3,19,20^ and motor^21^ populations in cortex exhibit varying levels of instability, raising the question of how perception and behavior remains stable^22^. One possibility is that a subset of neurons exhibit greater stability. We tested this by imaging mice with steady behavioral performance (**Figure S4**) over several weeks (**Table S1**; Methods). We imaged each mouse approximately once a week for 3 weeks using the same three 700-by-700 μm planes spaced 60 μm apart in depth.

In vS1, some touch neurons maintained their touch response across sessions whereas others changed (**Figure 5A**). Similar trends were observed for whisking and licking representations (**Figure S5**). The fraction of the various touch neurons remained stable over time, as did the fraction of whisking and licking neurons (**Figure 5B**). Representational drift can occur even when the responsive population size is stable, as neurons gradually transition in and out of the responsive population. Did the most responsive neurons exhibit lower levels of drift? To assess this, we measured the fraction of neurons that were part of a particular population on the first imaging day and remained so on subsequent days (**Figure 5C**). Among vS1 touch neurons, unidirectional single whisker and bidirectional single whisker neurons exhibited rapid turnover, with only 35.1 ± 11.2% and 51.5 ± 14.9% of the most responsive quartile of neurons that belonged to a given category remaining in that category a week later. For multiwhisker populations, however, 75.0 ± 4.1% of neurons in the most responsive quartile remained multiwhisker a week later, far more than the bottom quartile (33.2 ± 10.5%; top vs. bottom, p=0.016, Wilcoxon signed rank test, n=7 mice). Similar trends were observed for whisking neurons (top quartile remaining in-category after one week: 82.7 ± 9.0%; bottom: 45.2 ± 15.1%; top vs. bottom, p=0.016) as well as right-licking (top: 83.5 ± 7.7%; bottom: 44.2 ± 15.4%; p=0.016) and left-licking (top: 79.8 ± 14.5%; bottom: 32.7 ± 7.9%; p=0.016) neurons.

**Figure 5.**
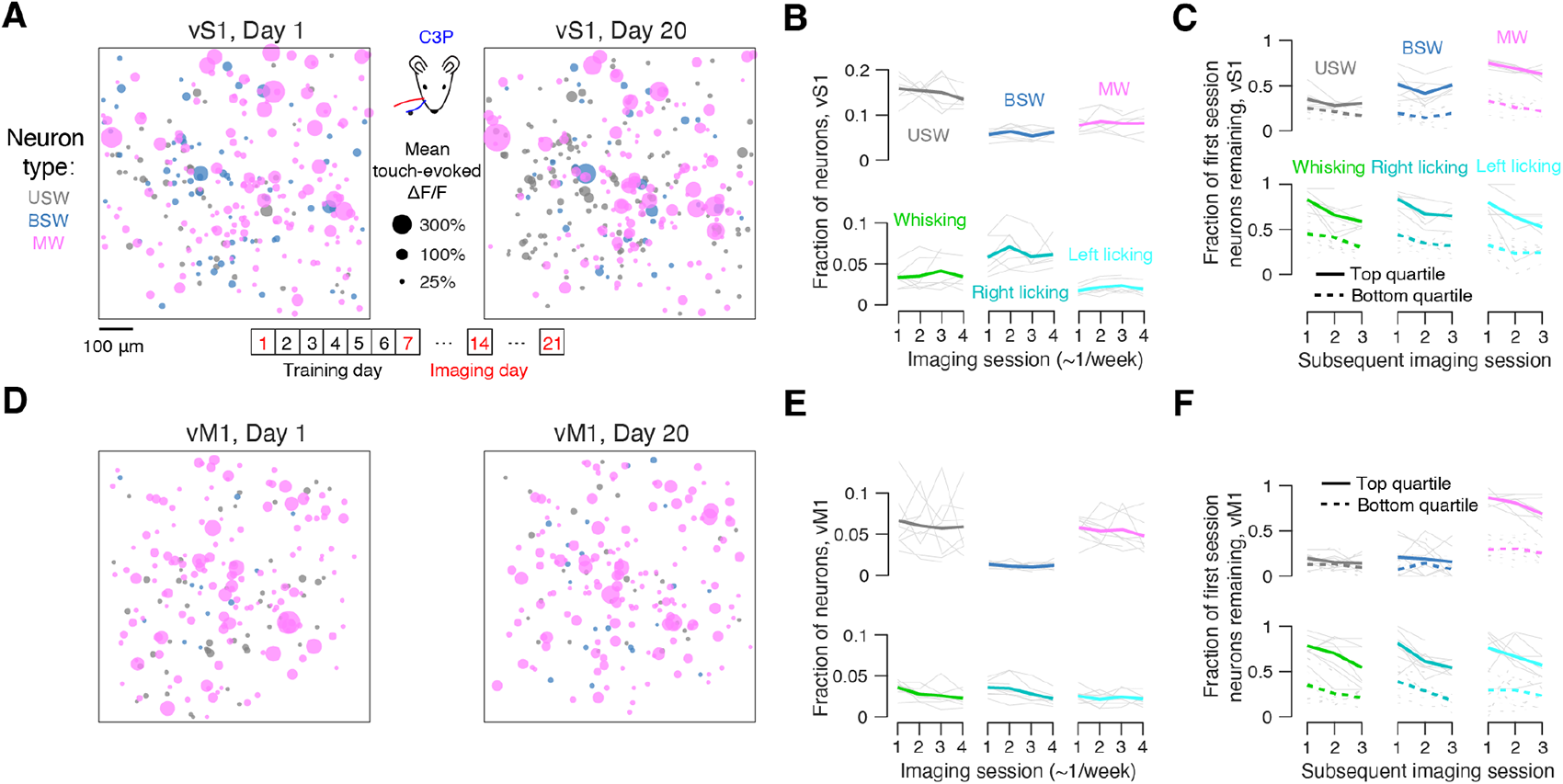
Dynamics of sensorimotor representations over several weeks. (A) C3 protraction touch-evoked ΔF/F responses collapsed across three imaging planes in an example mouse on the first and final imaging sessions (days 1 and 20). Circle size, mean touch-evoked ΔF/F. Bottom, typical timeline with four imaging timepoints in behaviorally stable animals. Specific representations are segregated by color. (B) Fraction of neurons belonging to a particular touch population across imaged sessions (∼3 weeks total, 1 week apart). Light gray lines, individual mice (n=7). Dark colored lines, mean. (C) Fraction of neurons that belonged to a particular population on day 1 that still belong to that representation on subsequent imaging sessions (week 1, 2 and 3 end). Data is sorted into top responsiveness quartile of a population (Methods; solid lines) and the bottom responsiveness quartile (dotted lines). Light gray lines, individual mice (n=7). Dark colored lines, mean. (D-F) As in A-C but for vM1.

Repeating this analysis in vM1, we similarly found that touch (**Figure 5D**) and movement-related (**Figure S5**) populations exhibited a mixture of stability and turnover. At the aggregate level, touch and movement populations were stable, with a constant fraction of neurons participating in specific populations (**Figure 5E**). As with vS1, even the most responsive quartile of narrowly tuned touch neurons had a low likelihood of remaining within-category a week later (**Figure 5F**; USW: 19.6 ± 9.7%; BSW: 20.7 ± 17.7%). Multiwhisker cells in the top responsiveness quartile, however, exhibited high levels of stability, with 86.7 ± 6.5% remaining multiwhisker cells after a week (bottom quartile, 29.6 ± 11.5%; top vs. bottom, p=0.004, Wilcoxon signed rank test, n=9 mice). Movement responsive populations were similarly stable (whisking top quartile remaining in-category after one week: 78.5 ± 12.8%; top vs. bottom, p=0.004; right licking top quartile remaining: 81.5 ± 17.3%; top vs. bottom, p=0.004; left licking top quartile remaining: 76.2 ± 22.4%; top vs. bottom, p=0.004).

Therefore, the top quartile of responders among touch, whisking, and licking neurons in both vS1 and vM1 exhibits greater longitudinal stability.

### Touch-motor interactions are mediated by the most responsive neurons

Neurons with mixed selectivity that respond to both touch and movement have been observed in both somatosensory^42,43^ and motor^5,44^ cortices. In vS1, we found neurons that responded to only touch, whisking, or licking, as well as neurons with mixed selectivity for touch-whisking and touch-licking (**Figure 6A**). Both single whisker and multiwhisker neurons overlapped with motor representations (**Figure 6B**). Mixed selectivity touch-whisking neurons were predominantly drawn from the multiwhisker population (**Figure 6C**; overall percentage of USW-whisking neurons, 0.6 ± 0.3%; BSW-whisking, 0.5 ± 0.3%; MW-whisking, 1.9 ± 0.8%; MW-whisking vs. USW-whisking, Wilcoxon signed rank test, p=0.016; MW-whisking vs. BSW-whisking, p=0.016). Mixed selectivity touch-licking neurons were also mostly drawn from the multiwhisker population (USW-right licking neurons, 1.1 ± 0.4%; BSW-right licking, 0.9 ± 0.3%; MW-right licking, 1.9 ± 0.7%; MW-right licking vs. USW-right licking, p=0.047; MW-right licking vs. BSW-right licking, p=0.016).

**Figure 6.**
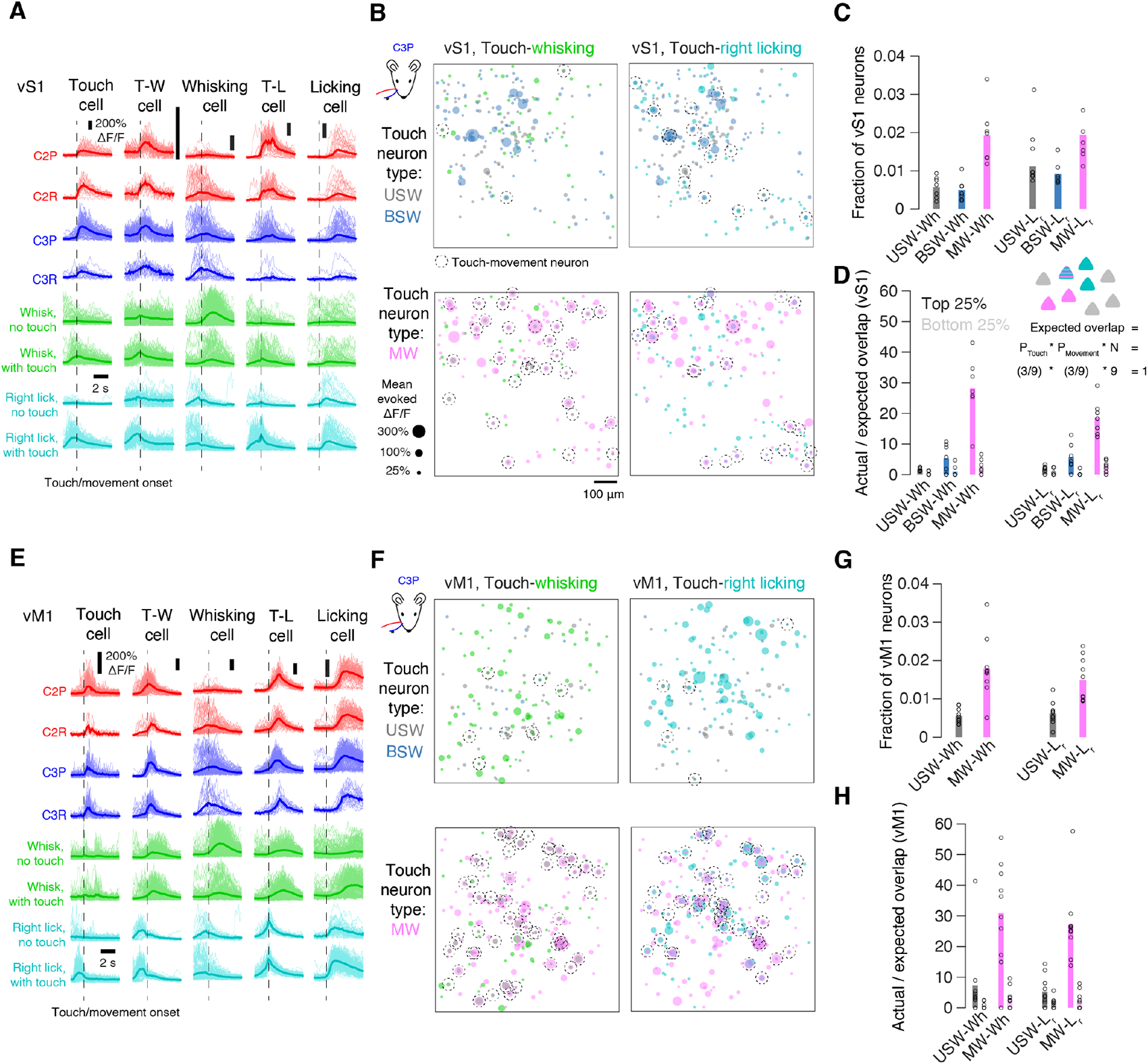
Multiwhisker touch neurons participate in whisking and licking representations. (A) Example vS1 cells. Each column shows the responses of a single cell aligned to touch or movement. Top to bottom, four basic touches; whisker movement-aligned responses on trials without and with touch; right licking-aligned responses on trials without and with touch. Left to right, touch cell; touch-whisking cell; whisking cell; touch-licking cell; licking cell. (B) Overlap between touch and movement-related populations, collapsed across three imaging planes from an example animal. Top row, narrowly tuned touch cells (USW, BSW). Bottom, multiwhisker cells. Left column, whisking; right column, right licking. Dot size indicates mean evoked ΔF/F response to touch or movement. Neurons responding to both touch and movement are marked with a dotted circle. (C) Fraction of all neurons that had mixed selectivity, separated by touch type. Wh, whisking, L_r_, right licking. (D) Ratio of actual to expected overlap in various touch-movement populations. In all cases, darker left bar is the overlap for the top 25% of responders from both populations, lighter right bar is for the bottom 25%. Bars indicate mean; circles, individual mice (n=7). Inset, calculating the expected overlap between two representations. The expected overlap is calculated under the assumption of two independent draws from the imaged population. (E, F) As in A, B, but for vM1. (G, H) As in C, D, but for vM1 (n=9).

Were strongly responsive multiwhisker neurons especially likely to participate in movement representations? To address this, we computed the number of mixed selectivity neurons expected if membership in one population was independent of the other (**Figure 6D**; Methods). We then computed the actual number of mixed selectivity neurons of a given touch-movement pair. Finally, we divided the number of observed mixed selectivity neurons by the number expected by chance to obtain a ratio of actual to expected overlap. We performed this calculation for neurons in both top and bottom responsiveness quartiles. Multiwhisker touch-whisking neurons in the top quartiles of touch and whisking responsiveness were observed 28.2 ± 10.4 more often than expected by chance (n=7 mice), more than the bottom quartile (2.5 ± 2.5; top vs. bottom, p=0.016, Wilcoxon signed rank test). Similarly, multiwhisker touch-right licking neurons in the top quartiles of touch and licking responsiveness were observed 18.5 ± 5.8 more often than expected by chance (n=7 mice), whereas bottom quartile neurons occurred 2.7 ± 1.9 more often than chance (top vs. bottom, p=0.016).

We observed a similar pattern in vM1: both touch-whisking and touch-licking neurons were present in vM1 (**Figure 6E**, **F**). Touch-whisking neurons were mostly drawn from the multiwhisker population (**Figure 6G**; percentage of USW-whisking neurons, 0.5 ± 0.2%; MW-whisking, 1.8 ± 0.8%; MW-whisking vs. USW-whisking, Wilcoxon signed rank test, p=0.002, n=9 mice), as were touch-licking neurons (USW-right licking neurons, 0.6 ± 0.3%; MW-right licking, 1.5 ± 0.6%; p=0.006). As in vS1, the ratio of actual to expected number of multiwhisker touch-whisking neurons was higher than expected by chance among the top quartile of responders (**Figure 6H**; 30.8 ± 16.8, n=9 mice), and less so for the bottom quartile of responders (2.9 ± 3.4; top vs. bottom, p=0.004, Wilcoxon signed rank test). Multiwhisker touch-licking overlap was also greater than expected by chance for the top quartile of responders (27.0 ± 11.9; bottom: 2.2 ± 2.9; top vs. bottom, p=0.002).

The most responsive multiwhisker touch neurons in both vS1 and vM1 can thus potentially contribute to both whisking and licking population activity via these highly responsive mixed selectivity neurons.

### Touch-licking and touch-whisking neurons exhibit distinct dynamics

To determine how the mixed selectivity neurons could potentially influence movement, we examined dynamics in specific neural populations more closely. For touch-licking interactions, we averaged ΔF/F across specific neural populations aligned to touch or rightward licking and computed a cross-animal mean response, focusing on rightward licking as this was the lick direction paired with touch on correct trials. Among multiwhisker touch cells that were not lick responsive, neural activity showed clear increases locked to the time of touch, but not to licking (**Figure 7A**). For vS1 and vM1 cells that were right lick but not touch responsive, touch did not evoke immediate responses but instead drove an increase in average ΔF/F 1-2 s after touch, as expected from the lag between touch and right licks on these trials (**Figure 7B**). Licking aligned responses in lick-only neurons showed ramping activity several hundred milliseconds prior to lick on both touch and non-touch trials. In mixed selectivity multiwhisker touch-licking neurons, activity increased sharply upon touch in both areas (**Figure 7C**). These neurons showed strong ramping activity up to the time of lick both when licking was preceded by touch and when it was not, and exhibited far larger responses than licking- or touch-only neurons. Moreover, upon lick onset, both licking-only and mixed selectivity neurons showed immediate declines in activity.

**Figure 7.**
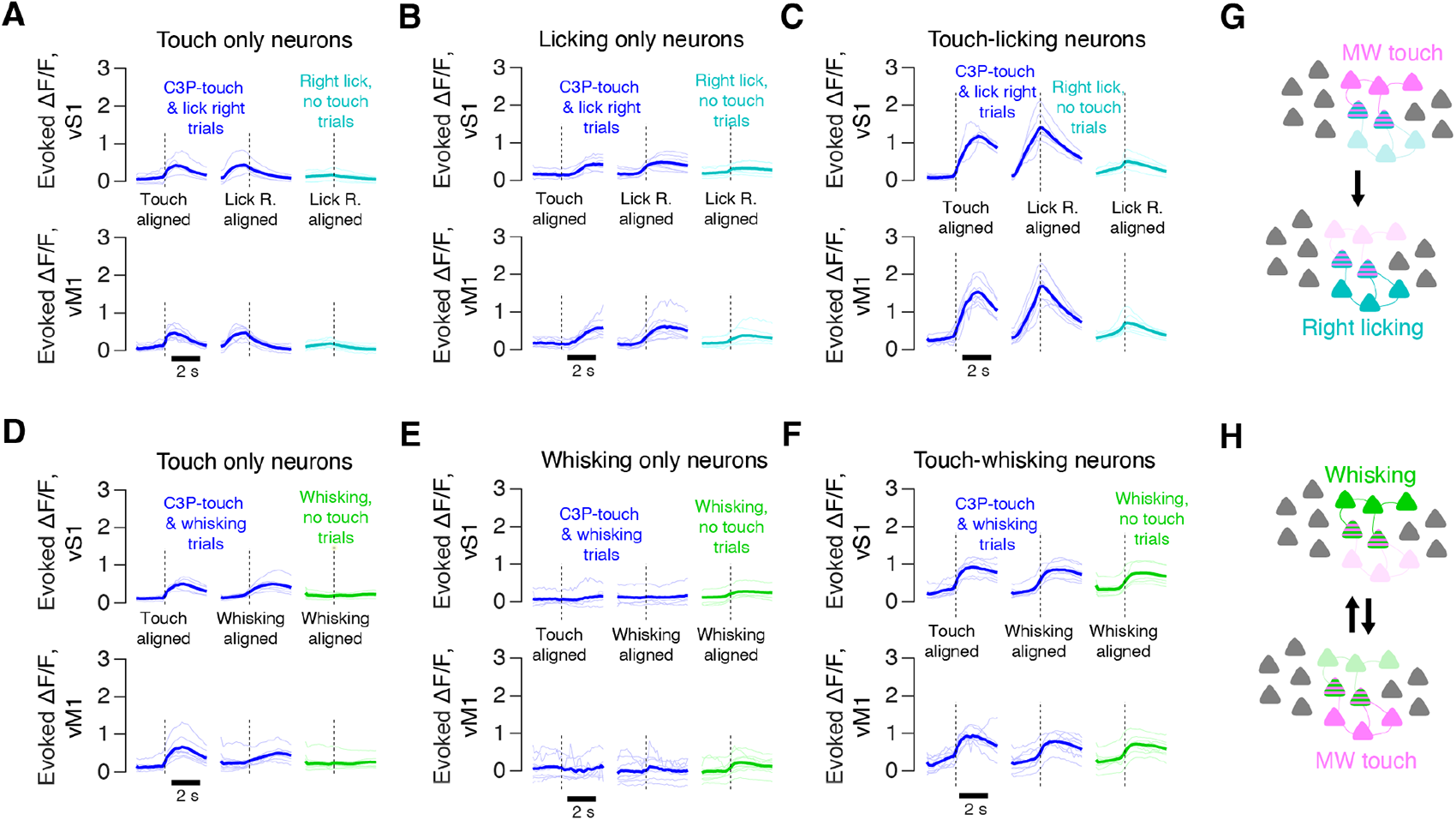
Touch-licking and touch-whisking neurons exhibit distinct dynamics. (A) Evoked ΔF/F response of multiwhisker touch-only neurons on C3 protraction touch trials (blue) and trials with licking but no touch (cyan). Touch trial responses are aligned either to the first touch (left; dotted vertical line), or lick onset (middle). Top, vS1; bottom, vM1. Thin lines, individual animals; thick lines, cross animal mean. (B) As in A, but for licking-only neurons. (C) As in A, but for touch-licking mixed selectivity neurons. (D) Evoked ΔF/F response of multiwhisker touch-only neurons on C3 protraction touch trials (blue) and trials with whisking but no touch (green). Touch trial responses are aligned either to the first touch (left; dotted vertical line), or whisking onset (middle). Top, vS1; bottom, vM1. Thin lines, individual animals; thick lines, cross animal mean. (E) As in D, but for whisking-only neurons. (F) As in D, but for touch-whisking mixed selectivity neurons. (G) Proposed circuit for transfer of activity from touch to licking populations via mixed selectivity neurons. Mixed selectivity neurons will be activated by touch, biasing the animal toward licking. (H) Proposed circuit for enhancement of multiwhisker touch responses during active whisking due to the engagement of mixed selectivity neurons. Mixed selectivity neurons will be activated by whisking, potentially strengthening touch responses during active whisking. Such a circuit could also function in the other direction, with touch activity driving greater whisking ensemble activation.

Did touch-whisking neurons behave similarly? As expected, multiwhisker touch cells that were not whisking responsive exhibited sharp ΔF/F increases at touch onset, and no responses on whisking-only trials in both areas (**Figure 7D**), whereas whisking neurons that were not touch responsive only responded on whisking onset (**Figure 7E**). Mixed selectivity multiwhisker touch and whisking cells exhibited robust responses both at touch onset and whisking onset (**Figure 7F**). As with touch-licking neurons, touch responses in touch-whisking neurons were larger than in touch-only neurons. Further, whisking responses in touch-whisking neurons were substantially larger than in whisking-only neurons. In contrast to touch-licking neurons, whisking onset did not result in immediate activity declines, suggesting that whisking and licking activity invoke distinct circuit mechanisms upon movement onset.

We conclude that relatively small populations of mixed selectivity neurons exhibit dynamics that make them ideal bridges between sensory and motor assemblies. These mixed selectivity neurons may integrate touch input that they in turn feed into motor assemblies, potentially contributing to both movement onset and movement modulation (**Figure 7G**). Such circuits could also enhance sensory responses during movement (**Figure 7H**).

## DISCUSSION

We examine sensorimotor integration in vM1 and vS1. We find sparse representations of both touch and movement, with ∼1-2% of cells often carrying a near-majority of either touch or movement-related activity, and touch activity concentrated among multiwhisker neurons (**Figures 2, 3**). These highly responsive touch, whisking, and licking neurons are particularly important for interareal communication (**Figure 4**) and exhibit elevated longitudinal stability (**Figure 5**). Highly responsive multiwhisker neurons interact with highly responsive whisking or licking neurons 20-30 times more frequently than expected by chance (**Figure 6**). Through combined touch and movement ensemble participation, these neurons exhibit temporal dynamics consistent with direct shaping of motor activity (**Figure 7**). Our work suggests that sensorimotor integration is mediated by sparse and specific populations of neurons with unusually strong responses, long-term stability, and elevated involvement in interareal communication.

Growing evidence suggests that cortex is organized into assemblies^56,57^ – groups of excitatory neurons that respond to the same stimulus and are interconnected^31–33,37^. Assemblies are ideally suited for integration of evidence in preparation for movement generation^58^. Touch-licking neurons may therefore directly contribute to licking behavior, as stimulating even a small number (<10) of licking-selective neurons can bias animal choice in a directional licking task^39^. Given the role of anterior lateral motor (ALM) cortex in licking^27^ and the progressive transfer of activity from vM1 to ALM^51^ in tasks similar to the one employed here, the touch-licking neurons observed in vM1 may directly bias animals toward licking (**Figure 7G**). Touch-whisking interactions are likely more complex: whisking activity decorrelates vS1 activity^59^ and likely increases sensitivity to external input. By directly activating touch-whisking neurons, whisking network activity may predispose the touch network towards stronger touch responses via pattern completion^34,35^ (**Figure 7H**). The converse may also be true: touch activity could influence whisking networks via touch-whisking neurons, potentially driving movement adjustments as object contacts are made^49^. Interestingly, upon licking initiation, we observe rapid silencing of the licking population, suggesting that unlike whisking, licking onset actively prevents touch responses. Targeted perturbation of mixed selectivity neurons using cellular resolution optical approaches^37,60,61^ will be needed to determine what role these neurons play in movement generation and modulation.

We find a high degree of similarity between vS1 and vM1. Both exhibit ultrasparse organization of touch, whisking, and licking. Both relay touch activity to one another predominantly via multiwhisker cells, with whisking and licking activity traveling in both directions. Multiwhisker touch and motor activity in both regions is surprisingly stable over several weeks, and interactions between multiwhisker touch and movement-related populations are found in both. The most prominent differences were the near absence of bidirectional single whisker cells in vM1, the lower number of unidirectional single whisker cells in vM1, and the higher number of right licking neurons in vS1. Responses in vM1 and vS1 did exhibit distinct dynamics within-category, with vM1 neurons far more likely to exhibit ramping activity prior to a movement, with some touch cells showing slow increases in responsiveness. Nevertheless, our results show that cortical sensorimotor integration is distributed across at least vS1 and vM1, with the populations involved actively communicating across these areas.

Neurons in sensory^3,19,20^ and motor^21^ cortices have responses that vary across days despite exposure to the same stimulus or execution of a stereotyped movement. This process is known as representational drift^22^. We find that the most responsive neurons exhibit greater stability compared to weakly responsive neurons (**Figure 5**). This suggests that drift can be high among the broader population while a smaller subset of neurons exhibits stability. The greater number of unstable unidirectional single whisker cells in vS1 means that, relative to vM1, the aggregate touch representation in vS1 will appear less stable, consistent with observations of stability in motor cortex and instability in sensory cortices.

Whisking activity in vS1 is known to be impacted by reafference^62^. It is likely that some subset of both whisking and licking cells observed here are not movement-generating but instead reflect reafferent sensory input. The temporal resolution of calcium imaging precludes using spike timing relative to movement to disambiguate these two possibilities. However, given that vS1 L4 selectively filters out whisker movement signals^63^, vS1 L2/3 whisker movement signals are more likely to originate from other cortical areas, such as vM1. Microstimulation experiments show that vM1, and specifically the vM1 touch patch, can trigger whisker movements^49,64,65^. Though it is possible that cortex only influences certain movements during early learning, even in well trained animals, cortex can bias movement, even if its removal does not impair movement production^66^. Presumably this reflects a broader set of both cortical and subcortical circuits that contribute to movement production. Nevertheless, touch-movement interactions in vS1 and vM1 can likely impact both whisking and licking.

Our work shows that touch, whisking, and licking populations are primarily driven by a small subset of highly responsive neurons. These neurons are far more likely to project between vM1 and vS1, exhibit elevated longitudinal stability, and are more likely to combine touch responses with either licking or whisking responses. We propose that sensorimotor integration in cortex is mediated by highly specialized populations of neurons acting as relays between ensembles processing sensation and those generating action.

## ACKNOWLEDGMENTS

We thank Alisha Ahmed, Andy Garcia and Lauren Ryan for comments on the manuscript. This work was supported by the Whitehall Foundation and the National Institutes of Health (R01NS117536).

## AUTHOR CONTRIBUTIONS

R.P. and S.P. designed the study. R.P. carried out most experiments with help from A.S. and M.L. R.P. and S.P. performed data analysis. R.P. and S.P. and wrote the paper.

## STAR METHODS

### Lead Contact

Further information and requests for resources and reagents should be directed to the lead contact, Simon Peron.

### Materials Availability

This study did not generate new unique reagents.

### Data and Code Availability

Source code used in this paper will be made available at http://github.com/peronlab upon publication. Data from this paper will be provided upon reasonable request to the authors.

## EXPERIMENTAL MODEL AND SUBJECT DETAILS

### Animals

We used adult Ai162 (JAX 031562) X Slc17a7-Cre (JAX X 023527)^54^ mice (8 female, 9 male; **Table S1**). These mice express GCaMP6s exclusively in excitatory cortical neurons. Both donor strains were bred and maintained as homozygotes, and homozygous parents were used for the cross. Breeders were fed a diet that included doxycycline (625 mg/kg doxycycline; Teklad) so that mice received doxycycline until they were weaned, suppressing transgene expression throughout development. All animal procedures and protocols were approved by New York University’s University Animal Welfare Committee.

## METHOD DETAILS

### Surgery

Mice 6-10 weeks old were implanted with cranial windows over the left hemisphere and headbars under isoflurane anesthesia (3% induction, 1-2% maintenance). A dental drill (Midwest Tradition, FG 1/4’ drill bit) was used to make a 3.5 mm diameter circular craniotomy in the left hemisphere over vS1 (center: 3.3 mm lateral, 1.7 mm posterior to bregma) or vM1 (center: 1.0 mm anterior, 0.8 mm lateral from bregma). A double-layer cranial window (most superficial: 4.5 mm diameter; closer to brain: 3.5 mm inner diameter, #1.5 coverslip; windows were adhered to each other with Norland 61 UV glue) was positioned over the craniotomy. For vM1 surgeries, bone wax (CP Medical) was applied to the bone overlaying the superior sagittal sinus to minimize bleeding. The headbar and window were both affixed to the skull using dental acrylic (Orthojet, Lang Dental). Mice were post-operatively injected with 1 mg/kg of buprenorphine SR and 5 mg/kg of ketoprofen.

### Retrograde labeling

Retrograde viral injections were performed in mice who had previously been implanted with a cranial window and imaged during behavior, or at the time of initial surgery. Previously imaged mice were first removed from water restriction for a week before retrograde viral injection. In all cases, the position of vascular features with respect to bregma was recorded on the initial surgery. Because the dental acrylic headcap covered vM1 in vS1 window animals, and vS1 in vM1 window animals, a small hole was drilled in the acrylic over either vS1 (3.3 mm lateral, 1.7 mm posterior to bregma) or vM1 (1.0 mm anterior, 0.8 mm lateral from bregma). After reaching the skull, a dimple was drilled that allowed for pipette penetration. In both cases, we injected 100 nL of pAAV-FLEX-tdTomato (Addgene, 28306-AAVrg, 1×10¹³ vg/mL diluted 1:50 in 1X PBS) into the target area at a depth of 300-350 μm and a rate of 20 nL/min (Narshige MO-10 hydraulic micromanipulator). Injection was performed using a glass capillary pulled with a micropipette puller (P-97, Sutter) and beveled to a tip with a ∼25**°** angle and 25 μm diameter. This was backfilled with mineral oil and 2 μL of the virus was pulled into the tip. The pipette was lowered into the target area at a rate of 300 μm/min which was followed by a 1-minute delay before injection began. A subset of retrogradely injected animals were perfused (described below) and imaged on a confocal microscope (model SP5, Leica) using a 20x objective (**Figure S3**).

### Behavior

After surgical recovery, mice were water restricted and placed on a reverse light cycle. Mice were typically given 1 mL of water per day with small adjustments made to keep weight at 80-90% of pre-restriction baseline. Mice were trimmed to whiskers C2 and C3 and subsequently trimmed every 2-3 days.

Water-restricted mice were habituated to the behavioral apparatus for 2 days by head fixing them for 15-30 minutes and giving them free water. Mice were then trained on a two-whisker touch detection task in which a pole was presented either next to the tip of C2 or C3, between C2 and C3, or completely out of reach (**Figure 1A**). Pole presentation was via vertical movement driven by a pneumatic actuator (Festo). The pole position on each trial was randomized, and all three proximal positions had a frequency of 16.7%. Pole positions were occasionally adjusted if animals changed their resting whisker position. For the longer whisker, pole placement was beyond the reach of the shorter whisker. Though the longer whisker did occasionally touch on trials meant for the shorter whisker, we obtained large numbers of isolated touch trials for all whiskers and touch directions (**Figure 1C**). The pole was presented to the animal for one second, followed by a one second delay, after which a response cue (3.4 kHz, 50 ms) signaled the start of a one second response period. Mice would lick during the response period and would receive water for licking right on in-reach positions and left on out-of-reach positions. A loud (60-70 dB) white noise sound was played for 50 ms following the onset of pole movement, which encouraged appropriately timed whisking. The lickport was moved along the anterior-posterior axis via a motor (Zaber) so that it was only accessible during the response period. Mice were trained on this task until they reached proficiency (>70% performance). Imaging was only performed in well-trained animals. Training took place on separate training rigs, and these were used for longitudinal experiments to maintain stable performance between imaging sessions.

A BPod state machine (Sanworks) and custom MATLAB software (MathWorks) running on a behavioral computer (System 76) controlled the behavioral task. Sounds were produced and controlled by an audio microcontroller (Bela). Three motorized actuators (Zaber) and an Arduino controlled lickport motion. Licks were detected via a custom detection circuit (Janelia Research Campus).

### Whisker videography

Whisker video was acquired with custom MATLAB software using a CMOS camera (Ace-Python 500, Basler) with a telecentric lens (TitanTL, Edmund Optics) running at 400 Hz with 640 x 352 pixel frames. The video was illuminated by a pulsed 940 nm LED (SL162, Advanced Illumination) synchronized with the camera (typical exposure and illumination duration: 200 μs). 7-9 s of each trial were recorded, which included 1s prior to pole movement, the pole in-reach period, and several seconds following pole withdrawal. Data was processed on NYU’s High Performance Computing (HPC) cluster. Whiskers were detected using the Janelia Whisker Tracker^55^. Whisker identity assignment was then refined and evaluated using custom MATLAB software^3,37^. Whisker curvature (κ) and angle (θ) were then calculated at specific locations along the whisker’s length. Change in curvature, Δκ, was measured relative to a resting baseline curvature which was calculated at each angle independently. This value was obtained during periods when the pole was out of reach. Automatic touch detection was then performed, and touch assignment was curated manually using a custom MATLAB user interface^3^. Protractions were assigned negative Δκ values by convention.

### Two-photon calcium imaging

A custom MIMMS two-photon microscope (Janelia Research Campus; http://openwiki.janelia.org/wiki/display/shareddesigns/MIMMS) with a 16X objective (Nikon) was used for cellular-resolution imaging. An 80 MHz titanium-sapphire femtosecond laser (Chameleon Ultra 2; Coherent) tuned to 940 nm was used, with powers out of the objective rarely exceeding 50 mW. The microscope included a Pockels cell (350-80-02, Conoptics), two galvanometer scanners (6SD11268, Cambridge Technology), a resonant scanner (6SC08KA040-02Y, Cambridge Technology), a 16x objective (N16XLWD-PF, Nikon), an emission filter for green fluorescence (FF01-510/84-30, Semrock), an emission filter for red fluorescence (FF01-650/60, Semrock), and two GaAsP PMTs (H10770PB-40, Hamamatsu) and two PMT shutters (VS14S1T1, Vincent Associates). A piezo (P-725KHDS; Physik Instrumente) was used for axial movement. Three imaging planes spanning 700-by-700 μm (512-by-512 pixels) and spaced at differential depths apart depending on the experiment were collected simultaneously at ∼7 Hz; we refer to this group of planes as a ‘subvolume’. For ‘basic’ and ‘longitudinal’ experiments, one such subvolume with each plane spanning 60 μm was employed. For ‘retrograde’ imaging, three subvolumes with planes spanning 20 μm were used. In both cases, we imaged 180 μm in depth, starting at the L1-L2 interface (i.e., depths of ∼100-280 μm were imaged). Data from the flyback frame, during which the objective rapidly moved back to the topmost position, were omitted.

Scanimage (version 2017; Vidrio Technologies) was used to collect all imaging data, and power was depth-adjusted in software using an exponential length constant of 250 μm. In retrograde labeled animals with multiple subvolumes, each subvolume was imaged for about 100 trials before moving to the next subvolume. After the first day of imaging, a motion-corrected mean image was created for each plane, which was then used as the reference image for any subsequent imaging days. For ‘retrograde’ experiments (**Figure 4**), 1-2 days of imaging were used to ensure at least 150 cells per neuron. For ‘longitudinal’ imaging experiments (**Figure 5**), one imaging session of the same three planes was performed approximately one week apart at four time points (three weeks in time, total). For ‘basic’ experiments (all other figures), analysis used the first day of longitudinal imaging data.

After acquisition, imaging data were processed on the NYU HPC cluster. First, image registration was completed for motion correction using a line-by-line registration algorithm^3^. Segmentation was performed on one session: neurons from the first day of imaging were detected using an automated algorithm based on template convolution that identified neuron centers, after which a neuron pixel assignment algorithm that detects annular ridges given a potential neuron center^67^ was used to identify the precise edges of the neuron. All pixels, including the nucleus, were used. This initial segmentation was manually curated, establishing a reference segmentation for each plane. On subsequent imaging days, the segmentation was algorithmically transferred to the new mean images for a given plane for that day^5^. After segmentation, ΔF/F computation and neuropil subtraction were performed. The neuropil-corrected ΔF/F trace was used for subsequent analyses.

In animals with projection labeling, neurons were classified as projecting if their mean tdTomato fluorescence exceeded a manually selected threshold. For each pixel on a plane, the cross-session mean tdTomato fluorescence was calculated. For any given neuron, its ‘redness’ was taken as the mean red fluorescence value for its constituent pixels in this mean image. A manual user interface that flagged neurons exceeding a threshold red fluorescence was used to find the threshold which appropriately partitioned tdTomato expressing neurons from non-expressing neurons in each animal.

### Histology

After several days of imaging, some animals were perfused with paraformaldehyde (4% in PBS) and postfixed overnight. A vibratome (Leica) was used to cut coronal sections 100-mm thick which were mounted on glass slides with Vectashield antifade mounting media containing DAPI (Vector Laboratories). These sections were imaged on a fluorescent light microscope (VS120, Olympus).

### Area identification

The locations of vS1 (including individual barrel locations) and vM1 were identified by measuring the GCaMP6s ΔF/F at coarse resolution (4x objective, Nikon; field of view, 2.2 x 2.2 mm) on the two-photon microscope while the C2 and C3 whiskers were deflected together with a pole (**Figure S1**). Imaging was performed for a single imaging plane at 28 Hz. This was done in awake mice not engaged in any task. Cellular resolution imaging was centered on either the C2 and C3 barrels in vS1, or the vM1 touch patch.

### Neural classification

A neuron was classified as responsive or non-responsive for a particular touch trial by comparing ΔF/F_baseline_, the mean ΔF/F for the 6 frames (0.85 s) preceding the first touch, to ΔF/F_post-touch_, the mean ΔF/F for the period between the first touch and two frames after the final (interframe interval, ∼143 ms). We then averaged the ΔF/F_post-touch_-ΔF/F_baseline_ response across all touches of a given type, using only trials where the touch was isolated (i.e., no other types of touch occurred). If this number exceeded 0.1 (10%), the neuron was considered responsive to that touch type. Touch neurons were classified on the basis of which touches they responded to. Neurons responding only to a single touch type (**Figure 1C**, **E**) were considered unidirectional single whisker (USW); neurons responding to both directions of touch for a single whisker were considered bidirectional single whisker (BSW); neurons responding to any combination of at least two touches involving both whiskers were considered multiwhisker (MW).

To compute a neuron’s mean touch-evoked ΔF/F, we incorporated the response to all touch types for which that neuron was significantly responsive. In all cases, touch-evoked ΔF/F was measured as ΔF/F_post-touch_-ΔF/F_baseline_ for a given trial. For a given touch type (C2P, C2R, C3P, and C3R), mean touch-evoked ΔF/F was calculated using trials where only that touch type occurred. For unidirectional single whisker cells, we used the mean touch-evoked ΔF/F for the single touch type it responded to. For bidirectional single whisker cells, we used the mean touch-evoked ΔF/F across the two directions for the whisker the cell responded to, weighing both touch types equally. Finally, for multiwhisker cells, we first calculated the mean touch-evoked ΔF/F for the touch types to which that cell responded, then took the mean across these values.

Whisking responsive neurons were classified in the same way as described for touch responsive cells, with the key difference of aligning responses based on whisking bout onset as opposed to touch. Whisking bouts were defined as periods where the amplitude of whisking derived from the Hilbert transform exceeded 10°. As before, ΔF/F_baseline_ and ΔF/F_post-whisking-onset_ were compared and a cell was considered whisker responsive overall if it produced a mean whisking onset aligned response of at least 10%. For whisking this classification was performed using trials with no touch. Licking responsive neurons were classified by measuring the relative change in ΔF/F locked to the time of first lick. Licking neurons were further classified on the basis of which lick direction they produced the larger response for – right or left. Because licking took place 1-2 s after touch in our task, we could effectively classify licking cells without excluding touch trials.

### Response fraction analysis

For analyses estimating the fraction of the touch response carried by a particular cell type, we calculated, for each animal, the response of each neuron to each type of touch that that neuron responded to. For each trial of a given touch type, we computed the evoked ΔF/F (i.e., ΔF/F_post-touch_-ΔF/F_baseline_) across every neuron. We could then ascribe the fraction of response carried by USW, BSW, MW and non-touch neurons for that trial. For any given touch type (C2P, C2R, C3P, and C3R), we computed the mean fraction of activity carried by a given type of neuron averaged across all trials having only that touch type on a session. We only used C2P and C3P touches for these analyses as these touch types were more numerous and consistent in terms of kinematics. An analogous approach was used for whisking and licking responses.

### Mixed selectivity neuron analysis

For any two populations A and B, we computed the expected number of neurons belonging to both populations assuming they were randomly related, *E*(*N_A and B_*) = *f_A_* · *f_B_* · *N_neruons_* where *N_neurons_* is the total number of neurons and *f_x_* is the fraction of neurons belonging to population X. To calculate the overlap between A and B relative to chance, *0_AB_* we divided the actual number of neurons in both representations (*N_A and B_*) by the expected number: 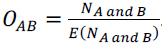.

### Quantification and statistical analysis

For comparisons between paired samples, we used the two-sided Wilcoxon signed rank test. For unpaired samples, we used the two-sided Wilcoxon rank sum test.

## SUPPLEMENTARY MATERIALS

**Figure S1. related to Figure 1.**
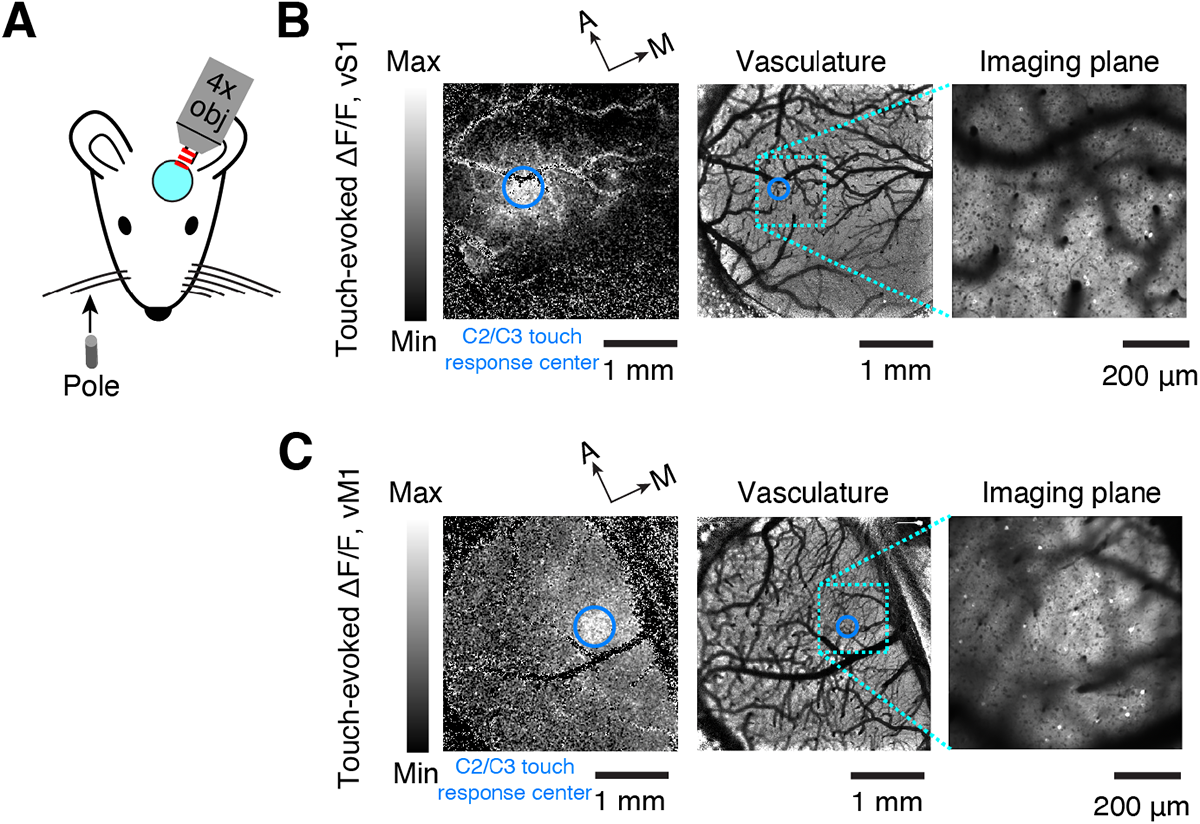
Identification of touch responsive regions in vM1, vS1. (A) Touch-responsive region identification procedure. The pole is moved against the two spared whiskers while most of the cranial window is imaged with a 4X objective (Methods). (B) Example animal touch-evoked response in vS1. Left, touch-evoked ΔF/F following pole touching C2 and C3. Middle, corresponding vasculature image with imaging plane outlined. Right, example imaging plane in vS1. (C) Same as B, but for vM1.

**Figure S2. related to Figure 2.**
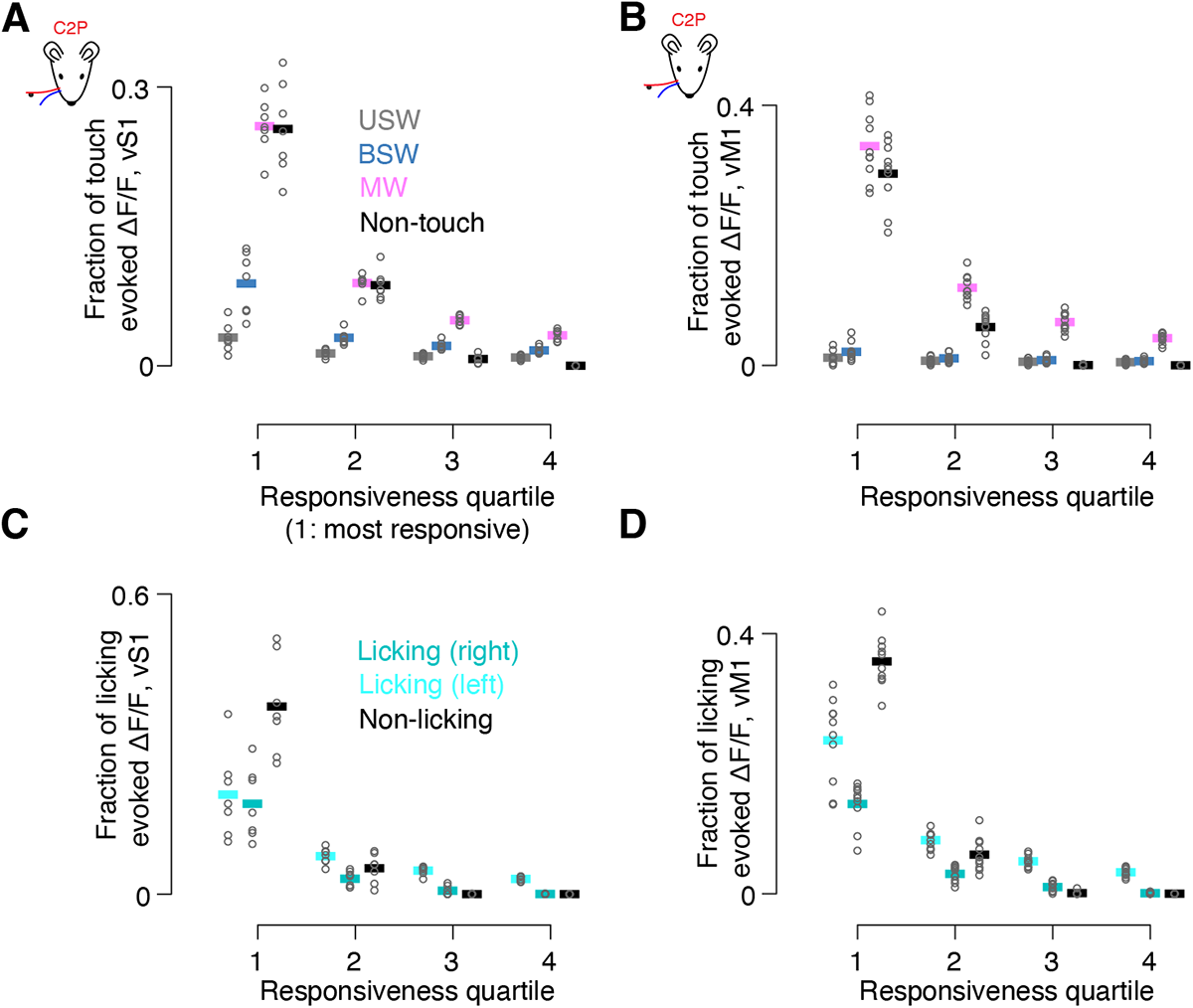
Concentration of sensorimotor responses for C2 protractions and left licks. (A) Fraction of touch-evoked ΔF/F contributed by each touch neuron type separated into quartiles by responsiveness (Methods). Only neurons that were considered responsive to C2P touches (Methods) were included. Left, most responsive 25% of neurons of a given neuron type. Thick line, cross-animal mean. Circles, individual animals. For an individual animal, all 16 values shown sum to 1. (B) Same A, but for vM1. (C) Fraction of left licking-evoked ΔF/F separated into quartiles by responsiveness (Methods). Thick line, cross-animal mean. Circles, individual animals. In all cases, for individual animals, all points sum to 1. (D) Same as C, but for vM1.

**Figure S3. related to Figure 4.**
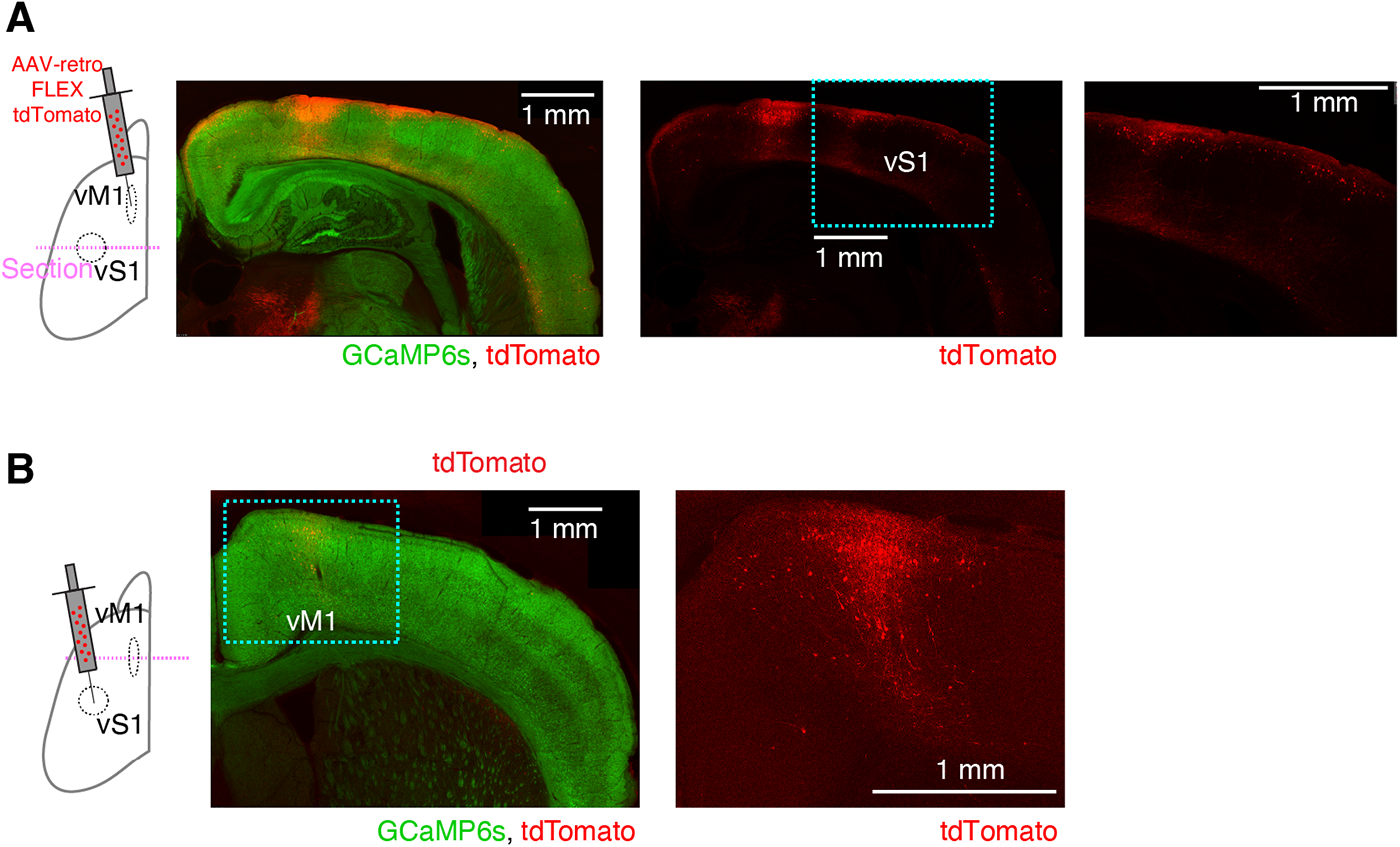
Confocal images of retrograde virus injected animals. (A) Confocal images of coronal section through vS1 in mouse with a retrograde AAV-FLEX-tdTomato injection in vM1. Left, dual color image including GCaMP6s. Right, tdTomato expression. In addition to vS1, more medial areas such including the dysgranular zone are labeled. (B) Confocal images of coronal section through vM1 in mouse with retrograde injection in vS1. Left, dual-color image. Right, tdTomato expression in vM1.

**Figure S4. related to Figure 5.**
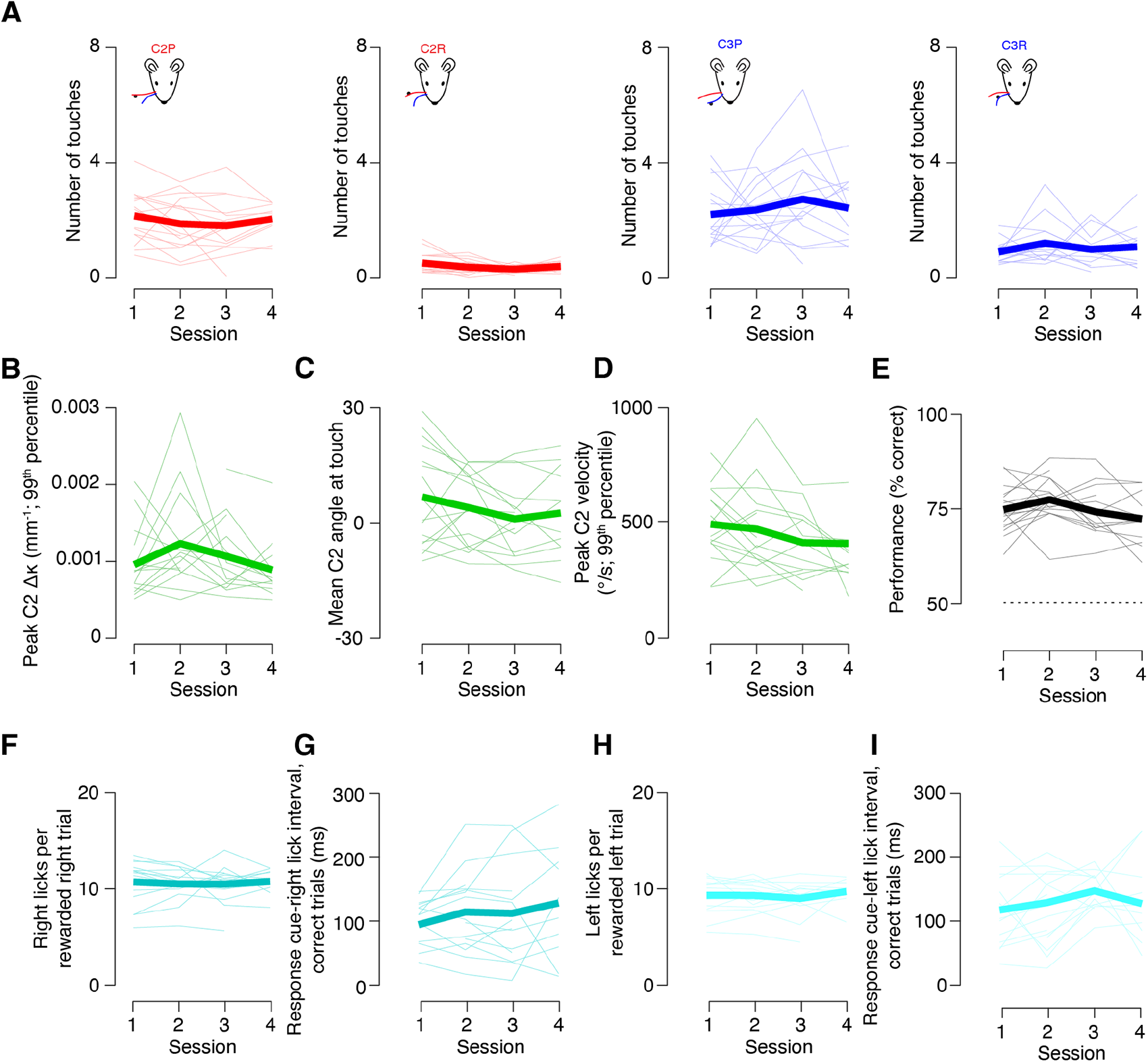
Behavior is stable during longitudinal imaging. (A) Number of touches across all imaged sessions and animals. Typically, imaging sessions were spaced one week apart. Left to right, C2P, C2R, C3P and C3R. Light lines, individual animals (n=15). Dark lines, mean across animals. (B) Peak change in curvature (Δκ) for C2 whisker across imaged sessions, quantified as 99^th^ percentile of Δκ values. (C) Mean angle at touch across all imaged sessions. (D) Peak velocity across all imaged sessions (**°**/s, 99^th^ percentile). (E) Task performance across all imaged sessions. (F) Right licks on correct trials (i.e., in-reach pole). (G) Time between response cue onset and first right lick on correct trials. (H, I) As in F, G but for left licks on out of reach trials.

**Figure S5. related to Figure 5.**
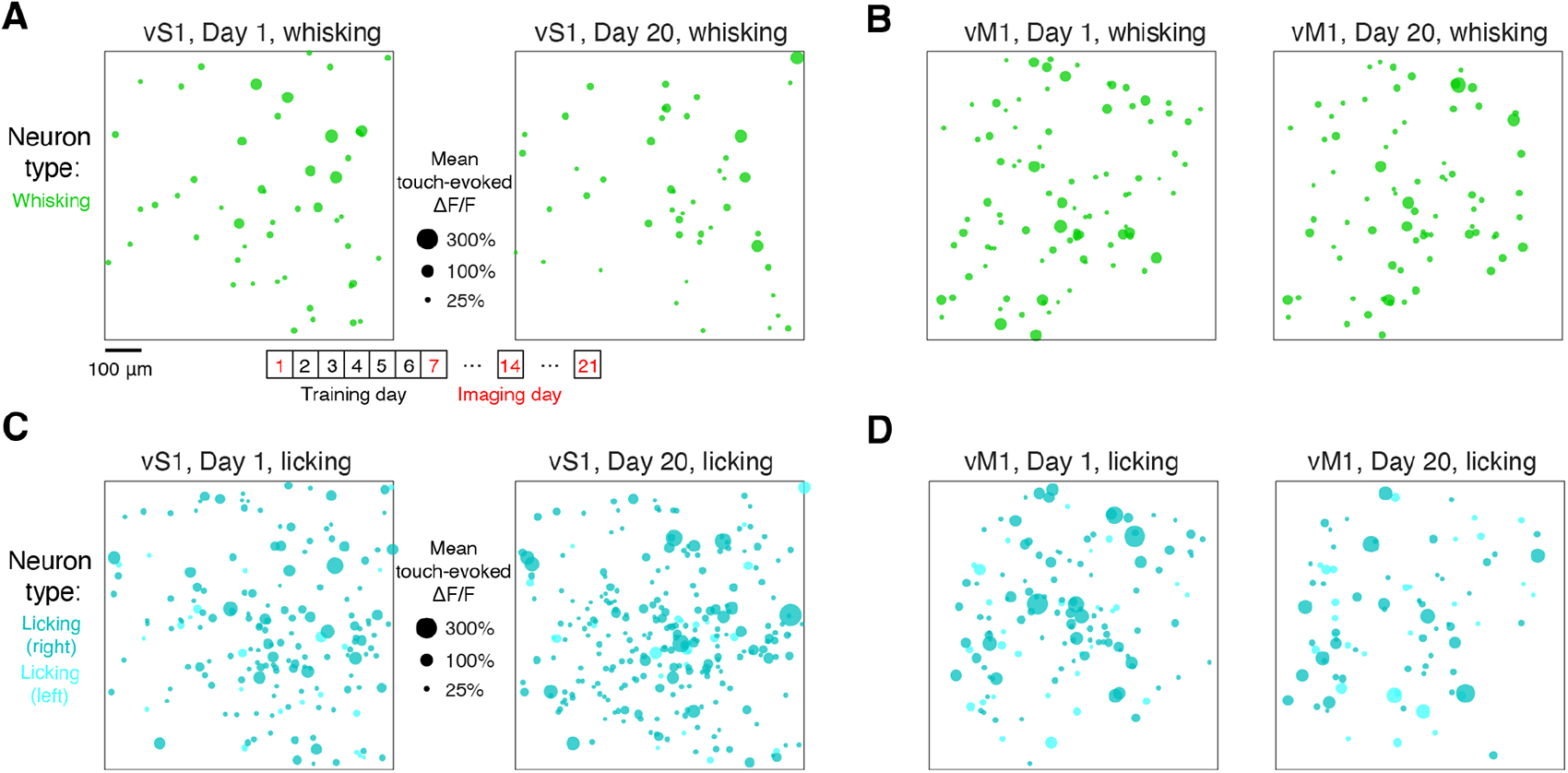
Longitudinal tracking of movement activity. (A) Whisking-evoked ΔF/F responses collapsed across three imaging planes in an example mouse on the first and final imaging session (days 1 and 20). Circle size, mean whisking-evoked ΔF/F. Bottom, typical timeline with four imaging timepoints in behaviorally stable animals. (B) As in A, but for vM1 animal. (C) Licking-evoked ΔF/F responses on right lick trials collapsed across three imaging planes in an example mouse on the first and final imaging session (days 1 and 20). Circle size, mean licking-evoked ΔF/F. Specific representations are segregated by color. (D) As in C, but for vM1 animal.

**Table S1 related to Figure 1.**
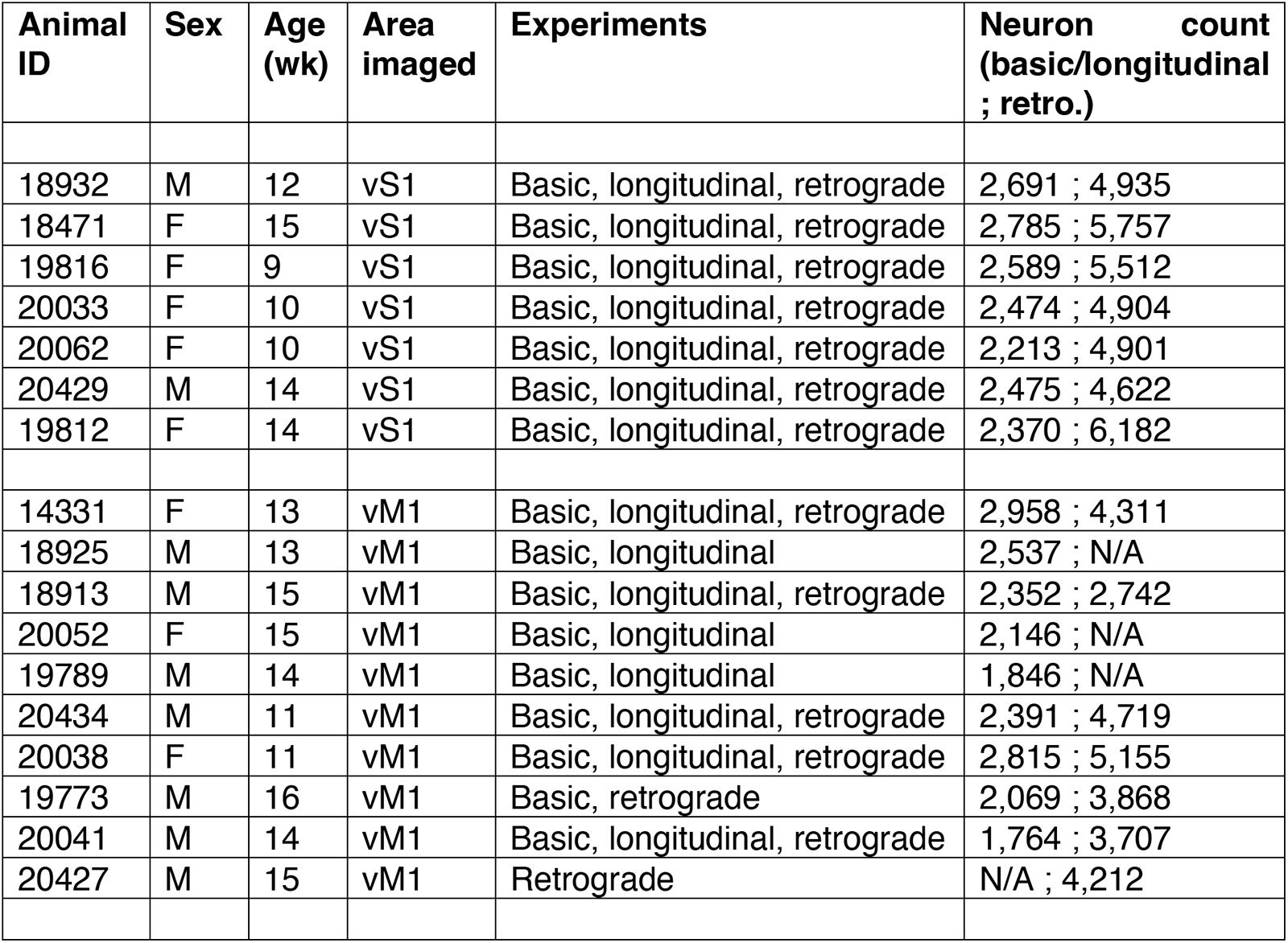
List of animals. All mice were transgenic Ai162 X Slc17a7-Cre, expressing GCaMP6s exclusively in excitatory neurons^54^. The C2/C3 whiskers were used in all animals. Age in weeks is provided for the first imaging day. Experiments include ‘basic’ analyses (**Figures 1**, **2**, **3**, **6**, **7**, **S2**), ‘longitudinal’ analyses (**Figures 5**, **S4**, **S5**), and ‘retrograde’ experiments that examine projecting neurons (**Figures 4**, **S3**). Cell counts are first given for basic and longitudinal experiments, which used the same imaging planes, and for retrograde experiments, which used different imaging planes to increase sampling density (Methods).

| Animal ID | Sex | Age (wk) | Area imaged | Experiments | Neuron count (basic/longitudinal ; retro.) |
| --- | --- | --- | --- | --- | --- |
| 18932 | M | 12 | vS1 | Basic, longitudinal, retrograde | 2,691 ; 4,935 |
| 18471 | F | 15 | vS1 | Basic, longitudinal, retrograde | 2,785 ; 5,757 |
| 19816 | F | 9 | vS1 | Basic, longitudinal, retrograde | 2,589 ; 5,512 |
| 20033 | F | 10 | vS1 | Basic, longitudinal, retrograde | 2,474 ; 4,904 |
| 20062 | F | 10 | vS1 | Basic, longitudinal, retrograde | 2,213 ; 4,901 |
| 20429 | M | 14 | vS1 | Basic, longitudinal, retrograde | 2,475 ; 4,622 |
| 19812 | F | 14 | vS1 | Basic, longitudinal, retrograde | 2,370 ; 6,182 |
| 14331 | F | 13 | vM1 | Basic, longitudinal, retrograde | 2,958 ; 4,311 |
| 18925 | M | 13 | vM1 | Basic, longitudinal | 2,537 ; N/A |
| 18913 | M | 15 | vM1 | Basic, longitudinal, retrograde | 2,352 ; 2,742 |
| 20052 | F | 15 | vM1 | Basic, longitudinal | 2,146 ; N/A |
| 19789 | M | 14 | vM1 | Basic, longitudinal | 1,846 ; N/A |
| 20434 | M | 11 | vM1 | Basic, longitudinal, retrograde | 2,391 ; 4,719 |
| 20038 | F | 11 | vM1 | Basic, longitudinal, retrograde | 2,815 ; 5,155 |
| 19773 | M | 16 | vM1 | Basic, retrograde | 2,069 ; 3,868 |
| 20041 | M | 14 | vM1 | Basic, longitudinal, retrograde | 1,764 ; 3,707 |
| 20427 | M | 15 | vM1 | Retrograde | N/A ; 4,212 |

